# A stable subgenomic reporter coronavirus enables transcriptional profiling of bystander cells

**DOI:** 10.64898/2026.02.27.708290

**Authors:** Ciaran Gilbride, Joe Hemsley-Taylor, Catarina Nunes, Rebecca Penn, James Boot, Nima Pieris, Rupa Tripathy, Ziyi Yang, Matthew Hutchinson, Olivia K Platt, Rachel Ulferts, Richard Mitter, Molly Strom, Nuno B. Santos, David L.V. Bauer, Harriet V. Mears

## Abstract

Insertion of fluorescent reporter genes into viral genomes is a powerful tool for monitoring infection. In coronaviruses, this is commonly achieved by replacing accessory open reading frames, thereby deleting endogenous gene functions. An alternative strategy is to manipulate viral RNA synthesis by inserting copies of the viral transcription regulatory sequence (TRS) which drive the transcription of viral subgenomic RNAs. However, coronavirus transcription is tightly regulated, and these modifications frequently disrupt native subgenomic RNA synthesis and attenuate viral growth. Here, we describe a reporter coronavirus that overcomes these limitations. Using human coronavirus (HCoV)-OC43 as a model system, we inserted an mNeonGreen reporter between the Spike and ORF5 coding regions, engineering the TRS and surrounding sequence to minimise off-target effects to transcription. This virus is genetically stable, with wildtype growth kinetics and unaltered subgenomic RNA transcriptional ratios. We developed a flexible reverse genetics system, which allows rapid cloning and virus recovery, supported by optimised HCoV-OC43 culture conditions, for high-titre stock generation, and validated analytical reagents. Our reporter virus enabled sensitive detection and isolation of infected cells, facilitating transcriptomic analyses that distinguish host responses in infected and bystander populations. We found that transcriptional responses to infection of cells in culture were predominantly inflammatory, rather than interferon-mediated, and that bystander cells upregulated pathways associated with cytokine response signalling and cell-cell contact sensing. Together, these tools expand the experimental utility of HCoV-OC43, an important seasonal respiratory pathogen and low containment model for betacoronavirus biology.

## Introduction

Coronaviruses are an ongoing threat to global health, with a large animal reservoir and high potential for zoonosis. In the last 25 years, three novel coronaviruses have emerged in the human population causing epidemics and pandemics of increasing global impact (1, 2). Limited spillover of canine coronaviruses in humans further highlights the persistent zoonotic risk posed by animal coronaviruses (3, 4). In addition to these emergent pathogens, four endemic human coronaviruses (HCoV) circulate worldwide and contribute to the global respiratory disease burden. Seasonal coronaviruses account for 15-30% of common cold cases, of which HCoV-OC43 is the most common (5-9).

While HCoV-OC43 infection typically results in self-limiting upper respiratory tract illness, disease can progress to pneumonia in immunocompromised patients, infants and the elderly (9-12). Rare cases of central nervous system infiltration have also been reported in mouse models and human patients, leading to viral encephalitis (13-16). HCoV-OC43 is a *Betacoronavirus*, in the same genus as the pandemic human coronaviruses, SARS-CoV, SARS-CoV-2 and MERS-CoV (Figure 1A). As such, HCoV-OC43 represents an appealing lower containment model for betacoronavirus biology.

**Figure 1.**
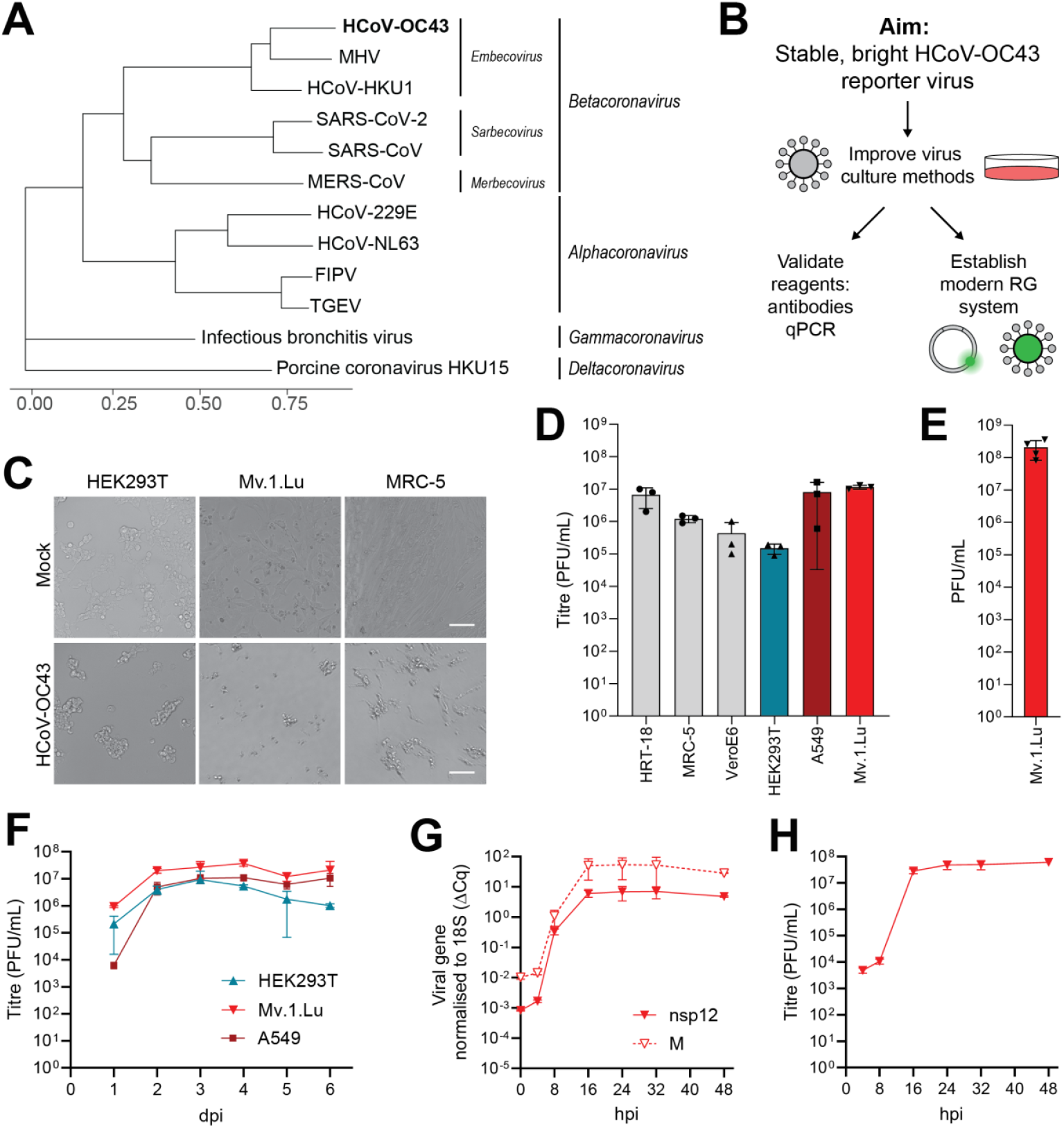
Cell culture conditions for high titre HCoV-OC43 growth. **A.** Phylogenetic tree of representative species from the family *Coronaviridae*, based on a codon-level alignment of open reading frame 1ab. **B**. Schematic outlining study aims. RG, reverse genetics. **C**. Light microscopy showing three cell lines (HEK293T, Mv.1.Lu, MRC-5) infected with HCoV-OC43, or mock infected, at an MOI of 0.0001, 5 days post infection (dpi). Scale bar represents 100 μm. **D**. Infectious titres of HCoV-OC43 released from the indicated cell lines at 5 dpi in plaque forming units (PFU) /mL. **E**. Infectious titres of HCoV-OC43 stocks from Mv.1.Lu cells. **F**. Growth of HCoV-OC43 in Mv.1.Lu, A549 and HEK293T cells infected at an MOI of 0.05. **G-H**. Growth of HCoV-OC43 in Mv.1.Lu cells infected at an MOI of 3, analysed by RT-qPCR (G) and plaque assay (H). Viral gene expression (nsp12, solid line and M, dashed line) was normalised to host 18S RNA (ΔCq). hpi, hours post infection. Data are means and standard deviations of at least three biological replicates.

Despite their importance, research into human coronaviruses has historically been constrained by poor growth in cell culture and a lack of effective analytical reagents. While HCoV-HKU1, another seasonal betacoronavirus, is refractory to culture in immortalised cell lines (17, 18), HCoV-OC43 can be grown to moderate titres – typically 10^6^ infectious units per mL of cell culture supernatant (19-21). Coronaviruses also possess exceptionally long RNA genomes (∼30 kb), often enriched with repetitive AU-rich regions, making them inherently difficult to manipulate using conventional molecular cloning tools (22-28).

Many modern reverse genetics systems rely on transformation-associated recombination (TAR) in *Saccharomyces cerevisiae*, to assemble cDNA fragments into stable yeast artificial chromosomes (29-32). These systems provide superior genetic stability over bacterial artificial chromosomes (25-27, 33), while allowing comparatively straightforward genetic manipulation (24, 34-36). Following the emergence of SARS-CoV-2, the development of reverse genetics systems for coronaviruses accelerated dramatically, with a focus on efficient, modular cloning strategies (37). Modern *in vitro* cDNA assembly methods, such as isothermal assembly, Golden Gate cloning, and circular polymerase extension reaction (CPER), have now been successfully applied to SARS-CoV-2 (38, 39) and other coronaviruses (40-43).

A common goal of viral reverse genetics is to engineer fluorescent or luminescent reporter viruses to monitor and quantify infection. In coronaviruses, tolerance to fluorescent tagging varies across species and genomic loci; for example, while some coronaviruses accommodate reporters at the 5′ end of the genome without substantial fitness costs, others exhibit attenuation (31, 44-46). Reporters are frequently inserted into the structural and accessory gene cassette in the 3′ end of the genome, typically replacing accessory proteins which are dispensable for replication in cell culture (46-50). However, to avoid deletion of viral genes, it is also possible to engineer dedicated reporter ORFs.

Coronavirus structural and accessory ORFs are expressed from a nested set of subgenomic RNAs (sgRNA), generated through a process called discontinuous transcription (51). During negative-strand synthesis, the viral RNA-dependent RNA polymerase complex encounters transcription regulatory sequences upstream of each ORF (TRS-B), which share short, conserved motifs that match the leader TRS (TRS-L) at the 5′ end of the genome. Polymerase template switching from a TRS-B to the TRS-L generates a shorter negative-strand sgRNA, which is subsequently copied into mRNA for the translation of that ORF. Artificial insertion of TRSs in the 3′ end of the genome can therefore drive the transcription of new sgRNAs (52-55) and generate fluorescent reporter viruses (30). However, the viral transcriptional programme can be disrupted in these viruses, resulting in attenuation and genetic instability (30, 54, 55).

To tackle these limitations, we developed a flexible reverse genetics system, using in vitro cDNA assembly, allowing reliable virus recovery within 8 days without the need for yeast recombination or in vitro transcription steps. We optimised cell culture conditions for HCoV-OC43, finding that mink lung cell lines support high titre growth, up to 10^8^ plaque-forming units per mL. We show that human embryonic kidney (HEK) 293T and lung epithelial (A549) cell lines support productive HCoV-OC43 replication, as well as validating analytical reagents and protocols, to expand the utility of HCoV-OC43 as a research model (Figure 1B).

Using this system, we generated a novel HCoV-OC43 reporter virus with a dedicated reporter sgRNA, which is genetically stable, with wildtype growth kinetics and an unaltered sgRNA transcriptional programme. Here, we demonstrate that this virus produces bright fluorescence, compared to insertion of the reporter in place of the NS2 accessory gene, enabling efficient sorting of infected and bystander cells. With this approach, we defined distinct transcriptional programmes in infected and bystander populations by RNA sequencing, that reflect coordinated inflammatory cytokine signalling. Together, our work represents a highly tractable toolkit for studying a human coronavirus infection at low containment.

## Results

### Cell culture models for high titre HCoV-OC43 growth

We first sought to identify a suitable cell line for high-yield HCoV-OC43 propagation, to support downstream reverse genetics development. We infected a panel of cells at a low multiplicity of infection (0.0001 PFU/mL) with HCoV-OC43 (ATCC Betacoronavirus 1, VR-1558). These consisted of MRC-5 human fibroblasts and HRT-18 human rectal tumour cells, which are frequently used for seasonal coronavirus culture, as well as VeroE6, which were recently reported as an optimal cell line for HCoV-OC43 growth (20). We included two further commonly used human cell lines, HEK293T and A549, as well as a mink lung cell line (Mv.1.Lu), which has previously been used to assess HCoV-OC43 growth by plaque assay (56).

After four days, we observed substantial cytopathic effect (CPE) in infected MRC-5, as expected, as well as CPE in both Mv.1.Lu cells and HEK293T cells (Figure 1C). When supernatants were analysed by plaque assay, we found high infectious titres from both Mv.1.Lu and A549 cells at five days post infection (Figure 1D). Stocks from Mv.1.Lu cells exceeded 10^8^ PFU/mL (Figure 1E) and amplicon sequencing showed no consensus-level deviations from the reference sequence. As such, Mv.1.Lu cells represent a new cell model for high titre HCoV-OC43 growth.

We then examined the kinetics of HCoV-OC43 growth in Mv.1.Lu cells, HEK293T and A549. Cells were infected at a low (0.05 PFU/cell) multiplicity and incubated for up to six days before supernatants were analysed by plaque assay and RT-qPCR. A primer-probe assay has been described previously (57), which binds within the M gene coding region. We used this as a basis to design a second primer-probe set which binds within nsp12, to measure genomic RNA replication. Melting temperatures were matched to the M primer-probes to facilitate multiplexing, and primer efficiency and specificity was verified using cDNA standard curves (Figure S1B-C).

We observed rapid replication in Mv.1.Lu cells, with high titres (>10^6^ PFU/mL) even at one day post infection, and peak titres at 2-4 days (Figure 1F). Titres from Mv.1.Lu cells were consistently higher than either A549 or HEK293T cells. Infectious titres correlated with genome replication, which increased rapidly between one- and two-days post infection, then waned (Figure S1D). We then infected Mv.1.Lu cells at a high (3 PFU/cell) multiplicity and monitored replication over 48 hours. Genome replication occurred between 4-16 hours post infection (Figure 1G), while infectious titres increased from 8-to-16 hours post infection, lagging behind genome replication by ∼4 hours (Figure 1H). Titres plateaued by 24 hours post infection, when substantial CPE was observed.

In parallel, we determined growth of two seasonal alphacoronaviruses, HCoV-229E and HCoV-NL63, in a wide panel of cell lines (Figure S2). We found that Mv.1.Lu cells did not support high titre growth of these viruses; rather Huh7.5.1 cells supported HCoV-229E growth (up to 10^7^ PFU/mL, Figure S2D), while LLC-MK2 cells supported low titre HCoV-NL63 growth (10^4^ PFU/mL, Figure S2E), consistent with standard protocols.

### Reagents for HCoV-OC43 analysis

In addition to our multiplexed qPCR assay, we evaluated reagents for immunodetection of HCoV-OC43 proteins. We tested two anti-N antibodies: a rabbit polyclonal antibody and a sheep-derived polyclonal antibody (48). Both antibodies detected N expression in Mv.1.Lu cells, following high multiplicity infection (3 PFU/cell), with robust detection at 16 and 24 hours, and as early as 8 hours post-infection for the sheep polyclonal antibody (Figure 2A). The sheep polyclonal also detected a second band with slower migration, consistent with phosphorylated N.

**Figure 2.**
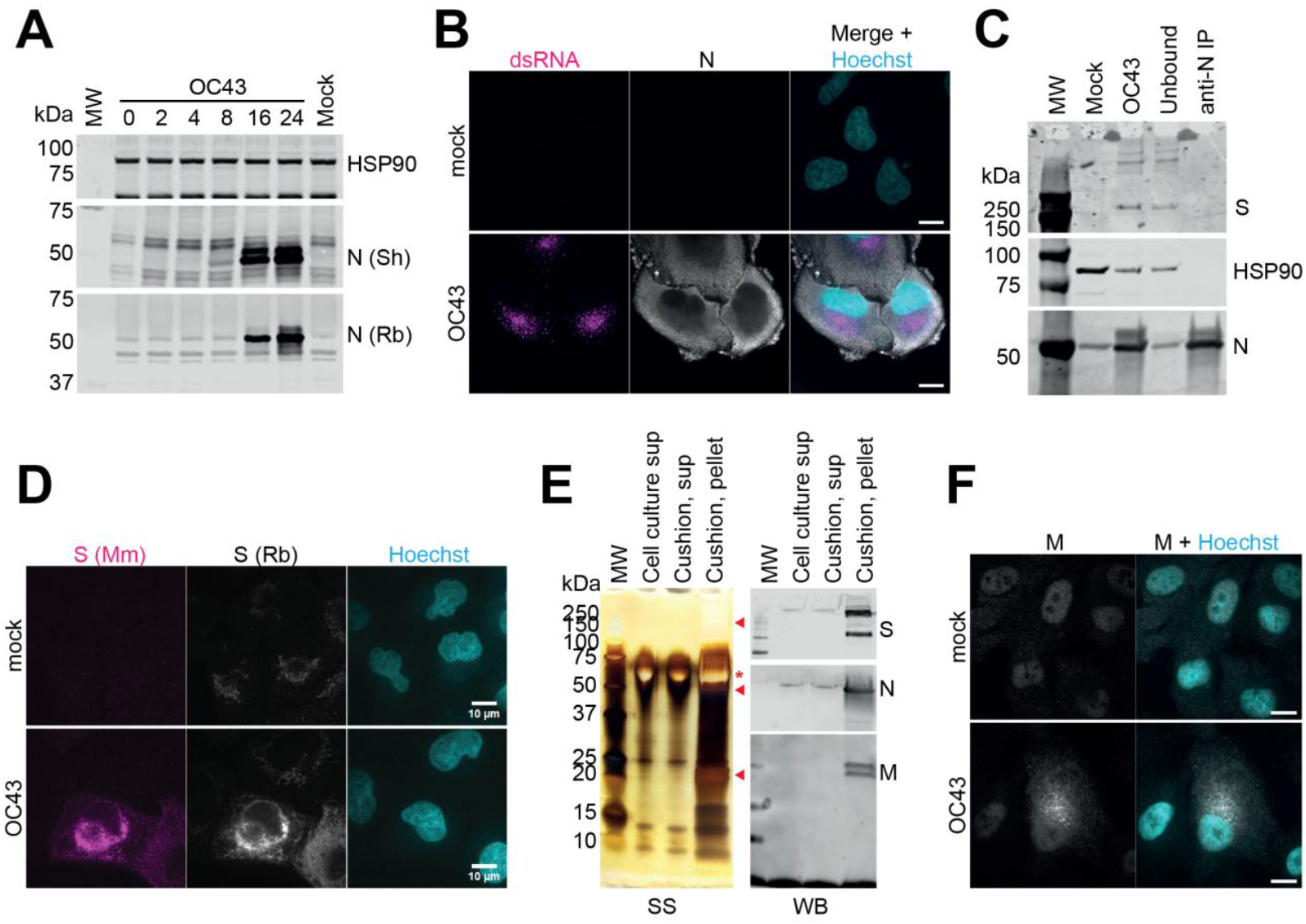
Reagents for analysing HCoV-OC43. **A.** Western blot of Mv.1.Lu cells infected with HCoV-OC43 at an MOI of 10 PFU/cell and harvested at the indicated time points, probed with two anti-Nucleocapsid (N) antibodies, derived from Sheep (Sh) polyclonal antiserum from the MRC PPU & CVR Coronavirus toolkit and a commercial Rabbit (Rb) polyclonal antiserum. Host HSP90 is included as a loading control. **B**. Immunofluorescence microscopy of HCoV-infected A549 cells, stained with Sheep anti-N and anti double-stranded RNA (dsRNA), a marker of RNA replication.**C**. Immunoprecipitation (IP) of lysates from HCoV-OC43-infected cells (MOI 3, 24 hpi) using Sheep anti-N and probed with Sheep anti-N and anti-Spike (S). Host HSP90 is included as a loading control. **D**. Immunofluorescence microscopy of HCoV-OC43-infected A549 cells, stained with mouse monoclonal (Mm) or rabbit polyclonal (Rb) anti-S. **E**. Silver stain (SS) and western blotting (WB) of virions purified by ultracentrifugation through a 30% sucrose cushion. The supernatant (sup) before and after sucrose cushion, and the pellet, containing purified virions, are shown. Viral structural proteins are indicated by red arrowheads and BSA from cell culture medium by a red asterisk. **F**. Immunofluorescence microscopy of HCoV-OC43-infected A549 cells stained with anti-SARS-CoV M. Scale bars represent 10 µm.

Immunofluorescence microscopy confirmed specific N staining with both antibodies, with minimal background in mock-infected controls (Figure 2B, Supplementary Figure S3A,B). Consistent with other coronavirus N proteins (48, 58), HCoV-OC43 N localised predominantly to the cytoplasm and cell periphery. Finally, immunoprecipitation using the sheep anti-N antibody successfully enriched N from infected cell lysates (Figure 2C). Spike protein was also detectable in infected cell lysates by immunoblotting (Figure 2C) and in virions purified by sucrose cushion ultracentrifugation (see Figure 2E) using a rabbit polyclonal antibody. By immunofluorescence microscopy, the rabbit polyclonal showed background punctate staining in mock-infected cells, consistent with mitochondria, while a mouse monoclonal antibody had low background in mock infection; both showed perinuclear staining in infected cells (Figure 2D), consistent with S accumulation in the ER.

No commercial antibodies specific to HCoV-OC43 Membrane (M) protein were available. Given the moderate sequence conservation across betacoronavirus M proteins, we hypothesised that some antibodies raised against heterologous M proteins might recognise HCoV-OC43 M. We therefore screened a panel of antibodies raised against M proteins from related betacoronaviruses for potential cross-reactivity. A monoclonal antibody against mouse hepatitis virus (MHV) M (59) failed to detect HCoV-OC43 M by immunoblot (Figure S3C). In contrast, a polyclonal antibody against SARS-CoV M weakly detected a band consistent with M in infected cell lysates at 24 hours post infection (Figure S3C). However, we noticed considerable background signal in mock-infected cells, precluding reliable detection of M in whole cell lysates (Figure S3D, lane 1). To reduce this background and enrich for viral structural proteins, virions were purified by sucrose cushion ultracentrifugation, which enabled definitive detection of M by western blot (Figure 2E, S3D, lane 4).

Finally, by immunofluorescence microscopy, the anti-SARS-CoV-M antibody displayed nonspecific nuclear staining in mock-infected controls, but additional punctate perinuclear staining specific to infected cells (Figure 2F), consistent with the localisation of other coronavirus M proteins (60-62). Together, these data indicate that, although background signal limits sensitivity, cross-reactive antibodies can be used to detect HCoV-OC43 M protein in purified virions and for imaging experiments.

### Generation of fluorescent reporter HCoV-OC43 viruses

We next developed a reverse genetics system for HCoV-OC43, leveraging modern *in vitro* assembly methods for rapid, high-fidelity viral assembly. The viral genome was divided into ten overlapping ≈3 kb fragments (F1-10), along with a terminal 1.5 kb fragment (F11) encompassing the genomic 3′ end (Figure 3A, B). A linker fragment (L) was designed to include a CMV promoter, to drive transcription of the viral genome in mammalian cells; for 3′ end processing, L encodes a poly-A tract and hepatitis delta virus ribozyme, followed by an SV40 polyadenylation signal and a Pol II transcriptional terminator (Figure 3A). Genome fragments were amplified from cDNA derived from HCoV-OC43-infected cells and cloned in pairs into modified pUC vector, termed pCLVR. To minimise bacterial toxicity associated with leaky expression of viral sequences, the cloning site in pCLVR is flanked by bi-directional *E. coli* transcriptional terminators (Figure S4A).

**Figure 3.**
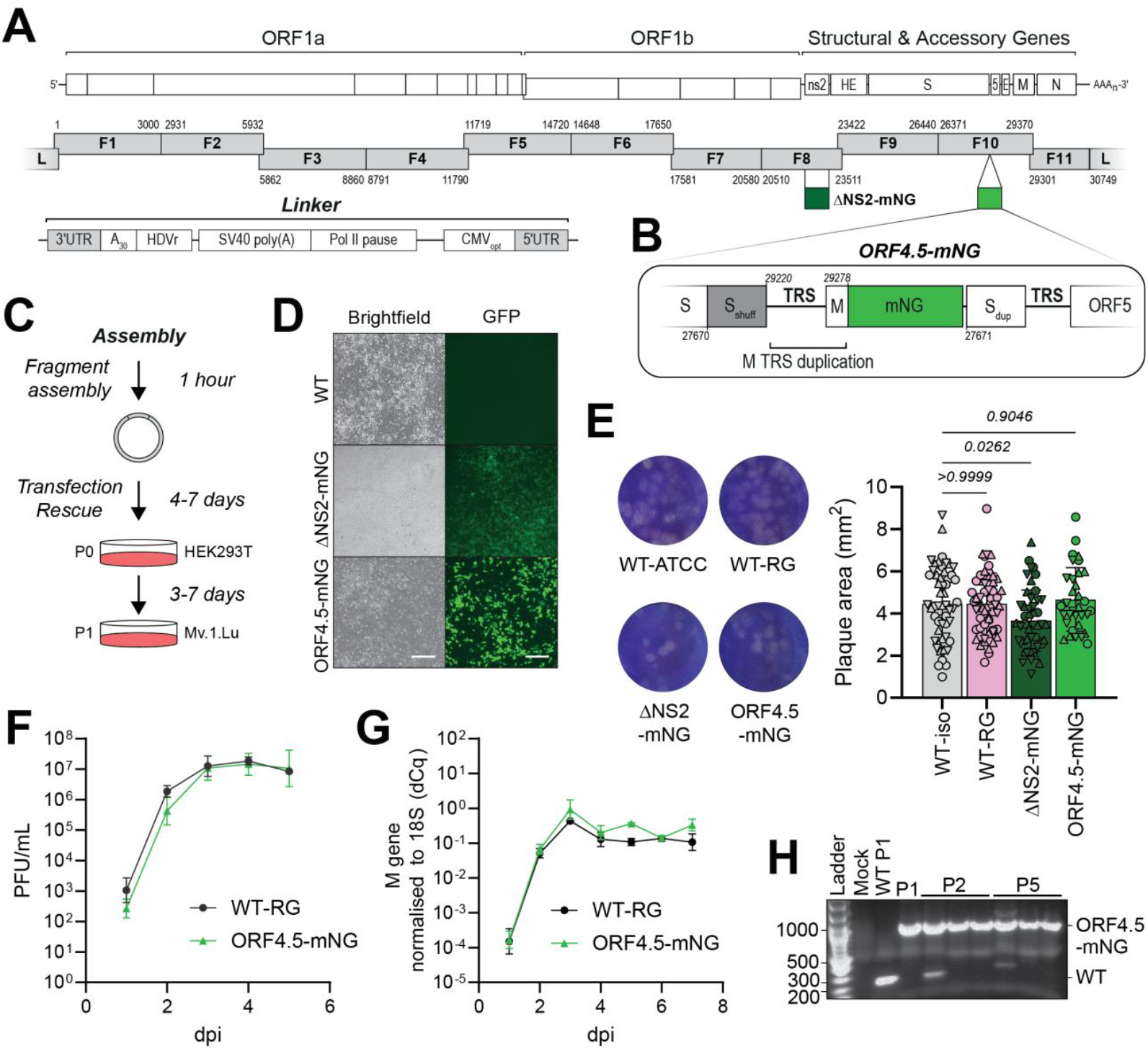
HCoV-OC43 reverse genetics (RG) by in vitro assembly from archived fragments. **A.** Schematic of the HCoV-OC43 genome, fragment and linker design, including a 5′ optimised CMV promotor (CMVopt) and 3′ 30-A tract, Hepatitis D virus ribozyme (HDVr), SV40 poly(A) signal and Pol II pause site. Insertion sites for fluorescent tags (mNeonGreen, mNG) are indicated. **B**. Details of the ORF4.5 insertion site. The 3′ end of the Spike (S) coding region was codon shuffled to prevent recombination (Sshuff). The M gene transcription regulatory sequence (TRS) and flanking sequence was inserted upstream of the mNG ORF; the genomic coordinates of the M TRS are indicated above. The wildtype 3′ end of the S gene was inserted downstream, to maintain the sequence context of the ORF5 TRS. **C**. Scheme for the assembly and rescue of infectious HCoV-OC43. **D**. Light and fluorescence microscopy showing Mv.1.Lu cells infected with RG-derived HCoV-OC43 viruses (MOI 0.0001, 4 dpi). Scale bars represent 200 μm. **E**. Plaque morphology of WT (ATCC isolate) and RG-derived HCoV-OC43 viruses (left) and quantification of plaque area (right). Data represent the means and standard deviations of three biological replicates, indicated by different symbols. Data were compared to WT-ATCC by one-way ANOVA and p values are shown. **F-G**. Growth of HCoV-OC43 in Mv.1.Lu cells (MOI 0.01), measured by plaque assay (F) and RT-qPCR against the viral M gene, normalised to host 18S rRNA (G). Data are means and standard deviations of four (F) or three (G) biological replicates.

This modular design results in a six-plasmid system that supports full-length genome assembly, to generate a single circular DNA containing the full-length HCoV-OC43 genome, with a 5′ promoter and 3′ polyadenylation and terminator elements. We found that circular polymerase extension reaction (CPER) assemblies were prone to precipitation and could not be stored prior to transfection, while isothermal assembly methods (e.g., Gibson or HiFi) were more stable and amenable to storage. Assembled genomes were transfected into HEK293T cells using liposomes and incubated for 4-7 days, until CPE was observed. Passage 0 supernatants were amplified on Mv.1.Lu cells for a further 3-7 days to generate a P1 stock (Figure 3C). For WT virus, titres of 10^8^ PFU/mL could be obtained within 10 days of initial assembly.

We designed two fluorescent reporter viruses (Figure 3A, B). First, we replaced the HCoV-OC43 NS2 open reading frame, with an mNeonGreen (mNG) reporter to create HCoV-OC43-ΔNS2-mNG. NS2 is a phosphodiesterase which antagonises the host antiviral factor RNaseL (63, 64) and is dispensable for replication in cell culture (65). The mNG coding sequence was codon-optimised to match the low GC content of the HCoV-OC43 genome. Fluorescence could be reliably observed at 4 days post-transfection, indicating successful rescue (Figure S4B); likewise, when Mv.1.Lu cells were infected at a low multiplicity of infection, the majority of cells showed green fluorescence at 4 days post infection (Figure 3D). HCoV-OC43-ΔNS2-mNG viruses displayed slightly delayed growth kinetics compared to WT virus (Figure S4C) and produced slightly smaller plaques (20% reduction in area relative to WT, Figure 3E), indicative of a minor loss of fitness. Nevertheless, after two (P2) and five (P5) passages in Mv.1.Lu cells, the mNG insert was maintained, determined by endpoint RT-PCR on RNA extracted from viral supernatants (Figure S4D).

To generate a reporter virus that preserves native viral protein function, we engineered a novel subgenomic RNA (sgRNA) to express mNG without disrupting endogenous ORFs. We modelled our synthetic TRS on the well-expressed M gene TRS, incorporating 28 nucleotides upstream and 24 downstream of the TRS core sequence (UCCAAAC), including predicted RNA structural elements. This region included the first 21 nucleotides of the M ORF, followed by the mNG reporter. To preserve the genomic context required for downstream TRS activity (i.e., ORF5), we appended the native 3′ end of the S gene after the mNG stop codon. To reduce the risk of homologous recombination between the duplicated S sequences, the upstream region was codon-shuffled while preserving the amino acid sequence. The resulting reporter sgRNA was designated ORF4.5 (Figure 3B).

HCoV-OC43-ORF4.5-mNG was rescued and grew to titres comparable to WT-RG virus, with similar growth kinetics (Figure 3F,G) and no changes to plaque phenotype (Figure 3E). We observed brighter fluorescence compared to HCoV-OC43-ΔNS2-mNG **H**. Agarose gel analysis of PCR amplicons surrounding the ORF4.5-mNG insertion site, amplified from RNA purified from supernatants, from passage 1 (P1), P2 or P5. Three biological replicates of P2 and P5 were analysed.

(Figure 3D), indicating the synthetic ORF is well expressed. Upon serial passage, the ORF4.5 insert was maintained up to P5 (Figure 4H) and fluorescent signal remained bright (Figure S4E,F). Together, these results show that the HCoV-OC43-ORF4.5-mNG is a stable, bright reporter virus with WT-like growth kinetics.

**Figure 4.**
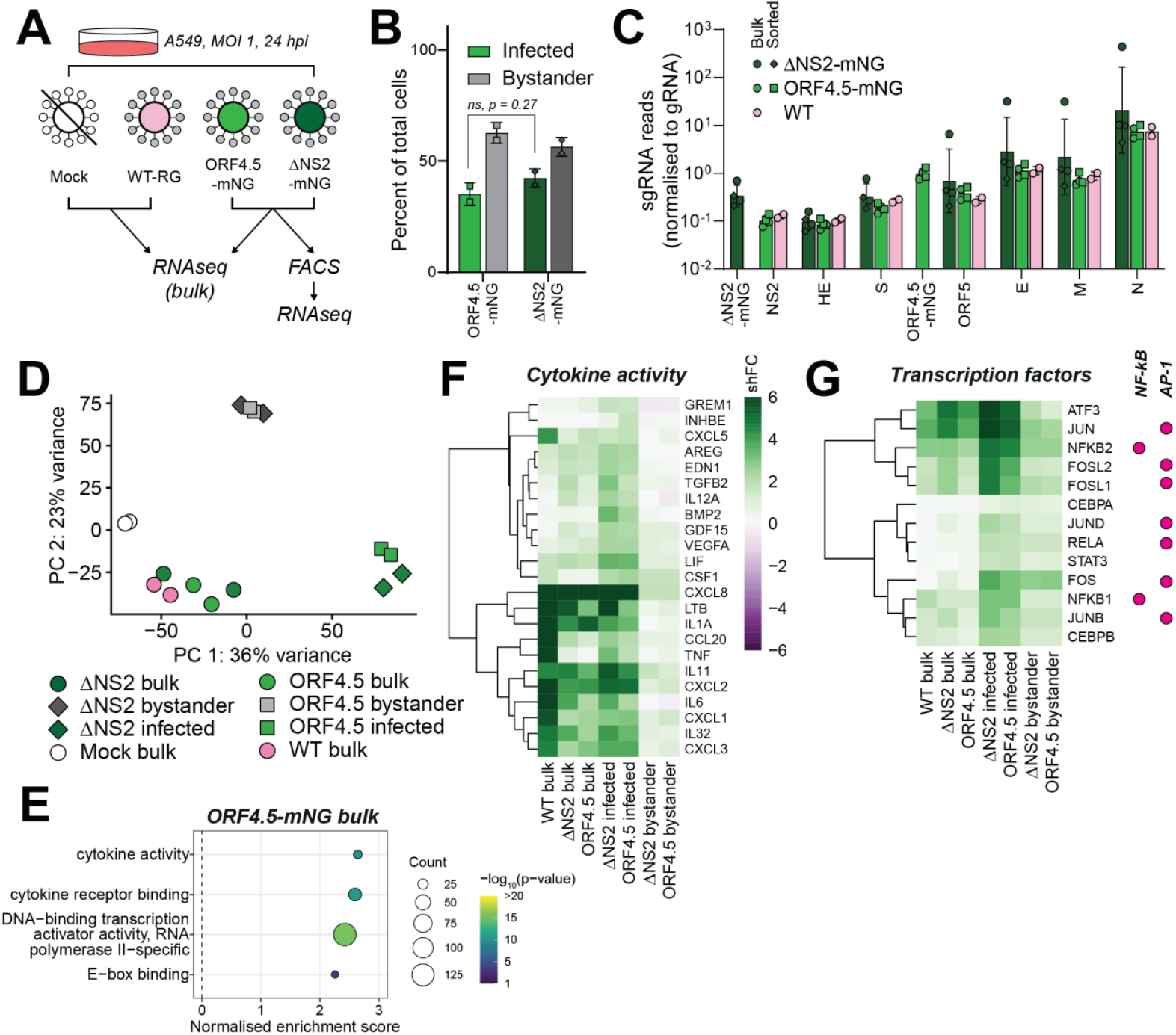
HCoV-OC43-ORF4.5-mNG transcription is comparable to wildtype and triggers an inflammatory cytokine response in infected A549 cells. **A.** Schematic showing the experimental plan for transcriptomic analysis. A549 cells were infected at an MOI of 1 and harvested after 24 hours. RNA was either extracted from the whole cell population (bulk) or cells were sorted by mNG fluorescence before RNA extraction from infected (mNG positive) or bystander (mNG negative) populations. **B**. Percentage of infected (green) and bystander (grey) cells after cell sorting. Infected cells were compared by Welch’s t-test. ns, not significant. **C**. Expression of viral subgenomic RNAs (sgRNA), determined by mapping chimaeric reads over the leader-TRS-body junction, normalised to genomic (gRNA) read counts. For each sgRNA, ratios for reporter viruses were compared to WT by one-way ANOVA and no significant differences were found. N.b. in DNS2-mNG, the NS2 TRS drives expression of mNG. **D**. Principal component (PC) analysis of transcriptomics data. **E**. Gene set enrichment analysis of differentially expressed genes in bulk ORF4.5-mNG-infected cells, based on molecular function. Functions are ranked by normalised enrichment score and coloured by significance. **F-G**. Heatmaps showing cytokine activity (F) and induction of pro-inflammatory transcription factors (G), coloured by shrunk log^2^ fold change (shFC) in expression compared to mock cells.

### Transcriptomic analysis of sorted infected and bystander cells

Finally, we aimed to understand how different populations of cells respond to infection, by profiling the host transcriptional response to HCoV-OC43 in both infected and bystander cells. Differentiating between infected and bystander cells usually relies on the detection of cell-surface viral antigens (66), which limits analysis to cells at later stages of infection. By contrast, our fluorescent reporters should facilitate sensitive detection and sorting of infected cells even early in the viral lifecycle, without the need for antibody staining. We infected A549 cells with HCoV-OC43-dNS2-mNG, HCoV-OC43-ORF4.5-mNG, or WT virus (Figure 4A). A549 cells are a human lung carcinoma cell line which can sense and respond to viral infection, and support productive HCoV-OC43 replication (Figure S5A). After 24 hours, RNA was extracted from the whole cell population (bulk) or following live-cell fluorescence-activated cell sorting.

We saw effective separation of infected (mNG-positive) and bystander (mNG-negative) cells for both reporter viruses (Figure 4B, S6). Consistently, we saw an enrichment of viral reads in infected cells following sorting, increasing from ∼50% of total RNA in bulk cell populations to ∼70% (Figure S6B). When we analysed viral transcription, we found that viral subgenomic RNA expression was comparable between our WT and reporter viruses (Figure 4C). Importantly, the insertion of ORF4.5 did not affect the transcription of up- or down-stream sgRNAs, while ORF4.5 itself was expressed to levels comparable to the M sgRNA. Likewise, we observed no consensus-level mutations in HCoV-OC43-ORF4.5-mNG, and a low frequency of single nucleotide polymorphisms, relative to the reference genome, supporting the genetic stability of this virus and the fidelity of our reverse genetics system (Figure S6C). Bystander cells showed 5-10% of reads mapping to the viral genome (Figure S6B). These cells may therefore represent dead-end infections or cells at an early stage of infection, before substantial viral protein production.

We then assessed host responses to infection. RNA-seq from bulk cell populations, infected with WT or fluorescent reporter viruses, showed highly similar transcriptional signatures, which were distinct from mock-infected controls, as shown by Principal Component Analysis (PCA, Figure 4D). In contrast, RNA-seq of FACS-sorted populations revealed that infected and bystander cells formed separate clusters, distinct from both mock and bulk samples, indicating that sorting effectively isolated transcriptionally unique cell populations (Figure 4D).

Analysis of bulk populations showed robust upregulation of inflammatory response genes following HCoV-OC43 infection (Figure 4E, F, Figure S7A-E). Infected cells displayed increased expression of multiple cytokines, including proinflammatory interleukins, and chemokines associated with neutrophil recruitment (Figure 4F). Indeed, among the most strongly upregulated transcripts in infected cells encoded for the neutrophil chemoattractant CXCL8 and the proinflammatory cytokines IL-1A and IL-6. We also observed induction of transcription factors, consistent with an early transcriptional response to cytokine signalling (Figure 4G). For example, we observe induction of NF-κB and AP-1 genes, which are typically induced in response to TNF and IL-1A. This may be counterbalanced by concomitant upregulation of anti-inflammatory factors such as TNFAIP3, which regulates NF-κB signalling to prevent excessive inflammation, and dual-specificity phosphatases, which control MAP-kinase activity (Figure S7G). In contrast, we did not observe a classical IFN response in infected cells (Figure S7H). Instead, we saw upregulation of TGF-β signalling (Figure S8A), which may contribute to suppression of IFN responses (67, 68).

Sorted, infected cells exhibited a substantially larger number of differentially expressed genes compared to sequencing the bulk cell population (compare Figure 5A and 5B, S7B and S7C). In addition to inflammatory signalling, we could also detect a greater degree of transcriptional remodelling (Figure 5C). In particular, there is upregulation of a subset of histone remodelling factors, transcriptional activators and transcriptional repressors, supporting selective and regulated reprogramming of transcriptional output.

**Figure 5.**
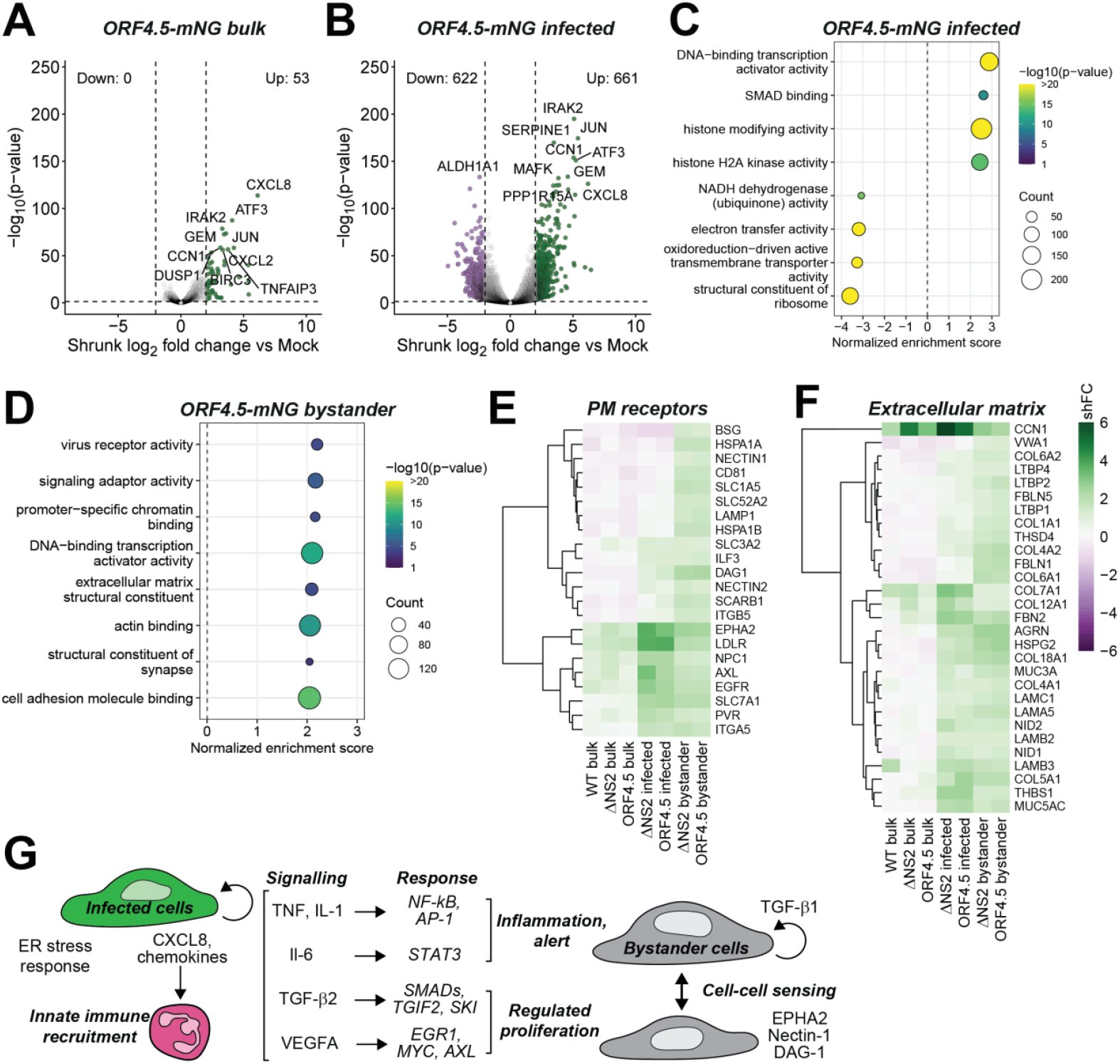
Sorting infected cells enriches for differentially expressed transcripts and reveals a unique transcriptional response in bystander cells. **A-B.** Volcano plots showing differentially expressed genes in HCoV-ORF4.5-mNG-infected cells, either from the bulk population (A) or after sorting (B), expressed as shrunk log^2^-fold change over mock infected cells. Significantly up-or downregulated genes (shrunk log2 fold change >±2, p value <0.05) are coloured and top 10 most significant genes are labelled. **C-D**. Gene set enrichment analysis of differentially expressed genes in sorted, ORF4.5-mNG-infected cells (C), or bystander cells (D), based on molecular function. Functions are ranked by normalised enrichment score (NES) and coloured by significance. **E-F**. Heatmap showing genes associated with plasma membrane (PM) receptors (E) and extracellular matrix structural components (F), coloured by shrunk log2 fold change (shFC) in expression compared to mock cells. **G**. Model for transcriptional responses in infected and bystander cells. Infected cells have a strong transcriptional response to HCoV-OC43, including secretion of pro-inflammatory cytokines and chemokines, as well as an ER stress response and downregulated metabolism. Bystander cells respond by upregulating pro-inflammatory pathways, including NF-κB, counterbalanced by mild anti-inflammatory and pro-proliferative signalling. Bystander cells upregulate factors associated with cell-cell sensing and wound repair.

Transcriptional responses in bystander cells were more muted than infected cells: although many genes were significantly upregulated (Figure S8B,C), the average fold change relative to mock cells was ≈2-fold, compared to ≈2.5-fold in infected cells (Figure S8D). In addition to genes associated with cytokine responsive pathways (Figure 4G, 5D, S8C), bystander cells upregulated a subset of TGF-β signalling genes which was distinct from infected cells, including both positive and negative regulators of TGF-β signalling (Figure S8A). In particular, infected cells showed higher expression the *TGFB2* isoform, while bystander cells preferentially expressed the *TGFB1* transcript, as well as transcripts encoding regulatory factors like SKI which represses TGF-β signalling. Bystander cells also displayed transcriptional changes consistent with alterations in plasma membrane composition and lipid-associated processes, including upregulation of genes encoding cell surface receptors and proteins involved in lipid metabolism that were not significantly induced in infected cells (Figure 5D, E). These included growth factor receptors and sensors of extracellular matrix disruption (Figure 5F). There was also upregulation of genes encoding actin cytoskeletal components, which was distinct from infected cells (Figure S8E). Together, this indicates sensing of local cell damage and cell migration in response to local damage.

Finally, we detected extensive downregulation of host transcripts in sorted, infected cells (Figure 5C), a feature largely absent in bulk data. These downregulated genes that were enriched encoded components of aerobic respiration and mitochondrial metabolism, consistent with a shift away from oxidative metabolism during HCoV-OC43 infection (Figure S9A). This was accompanied by upregulation of E-box-binding transcription factors, associated with metabolic and circadian rhythm regulation (Figure S9B). We also observed downregulation of genes encoding ribosomal proteins which, concomitant with an upregulation of genes encoding ubiquitin ligases involved in endoplasmic reticulum (ER)-associated degradation, such as *HRD1*, are indicative of ER stress and induction of the unfolded protein response (Figure S9C,D). Likewise, key effectors of the ER stress response were moderately upregulated in infected cells, including *CHOP* and *HERPUD1*. However, there was limited evidence of coordinated apoptosis induction at the transcriptional level downstream of the ER stress response. We observed modest changes to classical apoptosis markers, accompanied by the expression of anti-apoptotic genes such as *BIRC3* (Figure S9E). Together these data suggest that, while infected cells are experiencing translational stress, the response does not commit cells to apoptotic cell death.

Together, our data reveal distinct transcriptional changes in infected and bystander cells. Isolating infected cells enhanced signal-to-noise in transcriptomic analyses and revealed biological pathways that were obscured in bulk populations; likewise, analysis of bystander cells uncovered an early transcriptional response to the inflammatory cytokine milieu, characterised by programmes associated with tissue maintenance and repair.

## Discussion

In addition to its relevance as a human pathogen, HCoV-OC43 serves as an attractive, low-containment model for studying betacoronavirus biology and virus-host interactions. Nevertheless, its use has been constrained by suboptimal growth *in vitro* and a shortage of reliable reagents. Here, we report the development of a toolkit for studying HCoV-OC43, including an optimised cell culture model that supports high-titre viral replication, validated analytical reagents for qPCR and immunoassays, and a flexible, plasmid-based reverse genetics platform. These resources are provided in a detailed supplementary handbook. Using these tools, we engineered a fluorescent reporter virus, expressing mNeonGreen (mNG) from a dedicated subgenomic RNA, which was genetically stable and grew to wildtype titres.

To support efficient virus recovery and propagation, we first optimised virus culture conditions. HCoV-OC43 is frequently propagated in human fibroblasts (MRC-5), rectal tumour cells (HRT-18) (19-21) or, more recently, Vero E6 cells (20). We found that mink lung epithelial cells (Mv.1.Lu) supported robust HCoV-OC43 replication, with titres exceeding 10^8^ PFU/mL. Based on these findings, we established a two-step reverse genetics pipeline: rescue of recombinant virus in readily-transfected HEK293T cells, followed by amplification in Mv.1.Lu cells, enabling efficient production of high-titre recombinant HCoV-OC43 stocks. Sequencing stocks grown in Mv.1.Lu cells revealed no consensus-level mutations away from the reference sequence.

We used *in vitro* isothermal assembly to generate full-length HCoV-OC43 cDNA from overlapping fragments. Mutations and other genetic manipulations can be introduced into individual fragments by PCR, followed by a one-hour *in vitro* assembly reaction that is directly transfected into mammalian cells, enabling rescue of infectious virus within 4–7 days. Our system uses a CMV promoter to drive transcription of the viral genome in transfected cells, eliminating the need for an *in vitro* transcription step (69-71). This workflow substantially reduces the time required for virus generation compared with yeast-based assembly approaches, in which plasmid assembly and manipulation require ≈14 days, followed by an additional 12–16 days to recover infectious virus (72).

Although a cDNA BAC has been generated for HCoV-OC43 (16), we found that full-length cDNA was unstable and prone to deletion or recombination with bacterial DNA. This may be due to leaky expression of toxic viral gene products in *E. coli*, which hampers plasmid stability and can prevent successful propagation of full-length clones (22, 25). To mitigate this, we included bidirectional transcriptional terminators in our storage plasmid, pCLVR, which allowed HCoV-OC43 genome fragments to be stably propagated. Likewise, both *in vitro* ligation and circular polymerase extension PCR (CPER) were recently reported for HCoV-OC43 reverse genetics (73, 74). We found that CPER-assembled products were prone to precipitation, compromising rescue efficiency; as such, isothermal assembly offers a more flexible and reliable system.

We first replaced the NS2 accessory gene with an mNG reporter, which was codon optimised to match the low GC content of the HCoV-OC43 genome. Codon-optimisation is known to aid stability of coronavirus genetic insertions (47). NS2 is dispensable for MHV replication in cell culture (65) and has previously served as a site for reporter gene insertion in HCoV-OC43 reverse genetics systems (32, 69, 73, 74). Consistently, we found mNG was well tolerated at this site, with comparable peak titres and no evidence of reporter deletion after five passages in Mv.1.Lu cells, though we observed a slight delay in replication and a marginally smaller plaque phenotype, indicative of a small loss of fitness.

We then developed a reporter virus that preserves all endogenous ORFs, wherein the reporter gene is expressed from a dedicated subgenomic RNA. Insertion of viral transcription regulatory sequences (TRSs) can drive expression of novel subgenomic RNAs (54, 55, 75-77). Indeed, a recently described yeast-based HCoV-OC43 reverse genetics system inserted an artificial sgRNA between the M and N genes, using a minimal TRS core sequence to direct sgRNA transcription, to express a fluorescent reporter (30). However, reporter insertion toward the 3′ end can alter TRS usage and skew sgRNA expression profiles, reducing viral fitness (52-54). Consistently, Duguay *et al*. reported attenuation and altered transcriptional dynamics in multiple cell types (30).

We engineered our reporter, designated ORF4.5, between Spike and ORF5. Transcription was driven by a copy of the M gene TRS. Since coronavirus transcription efficiency is influenced by the RNA sequence context and local structures (51, 77-80), we preserved the surrounding sequence around both the inserted TRS and the downstream ORF5 TRS, to promote native transcriptional activity (79, 80). Our transcriptomic analysis demonstrated expression of native sgRNAs in the ORF4.5 reporter virus remained comparable to wildtype. The ORF4.5 insertion was also well tolerated, with little evidence of attenuation or deletion after serial passage, suggesting that the positioning of the reporter gene and preservation of native TRS flanking sequence elements is important for maintaining optimal viral fitness.

We then used these reporter viruses to investigate host responses to HCoV-OC43 infection in a human lung cell line. In bulk cell populations, we observed upregulation of proinflammatory cytokines but limited induction of canonical type I IFN response genes. This inflammatory response to infection, in favour of type I IFN gene expression, has previously been reported in MRC-5 cells infected with HCoV-OC43 (81) and A549 cells infected with SARS-CoV-2 (66, 82). Comparison of our recombinant viruses, with and without the NS2 protein, indicated that NS2 does not broadly alter innate immune signalling, consistent with its described role as a specific antagonist of RNaseL activation (64, 83-85). In addition to inflammatory signalling, we observed induction of ER stress response genes, a common feature of coronavirus infection (81, 86, 87), reflecting both the considerable translational stress during infection and the extensive remodelling of the ER during the establishment of viral replication-transcription complexes (88).

However, bulk RNA-seq has limitations for interrogating heterogeneous cell populations, including viral infections, since it samples an average across cells in different infection states (89). One solution is to perform single-cell RNA sequencing; however, this approach is costly and sequencing depth may be limited compared to traditional transcriptomic approaches, particularly for low abundance transcripts (90). Moreover, most high-throughput single-cell RNA-seq platforms exhibit a strong bias toward sequencing transcript termini (90, 91), which would limit the viral transcription analyses we performed here, where detection of sgRNAs relies on chimeric reads spanning leader-TRS-body junctions.

To overcome this, we sorted infected cells based on reporter fluorescence, to enrich for infected cells. Compared to our bulk RNA-seq results, we observed stronger upregulation of inflammatory and ER stress pathways, and enhanced our resolution of transcriptional reprogramming within infected cells. In particular, expression of genes encoding TNF and IL-1 family cytokines in infected cells correlated with the induction of NF-κB and AP-1 transcription factors, which was accompanied by the induction of transcriptional regulators and histone remodelling factors. Sorting infected cells also revealed a large number of downregulated transcripts, which were masked in bulk populations. These included ribosomal proteins, consistent with an ER stress response, as well as metabolic genes.

Finally, we examined transcriptional responses in bystander cells, which were stringently sorted based on low mNG reporter fluorescence, reflecting an absence of detectable viral translation. Consistently, bystander cells showed minimal induction of ER stress response genes or inflammatory cytokines. However, low levels of viral genomic RNA were detected in this population, suggesting that some bystander cells may have been recently infected or experienced non-productive infections. A previous report, which used cell surface expression of the viral Spike protein to differentiate infected and bystander cells in SARS-CoV-2 infection, similarly observed low levels of viral reads in bystander cells (66). However, the authors also observed that the transcriptional programme of bystander cells closely resembled that of mock-infected cells, with the exception of a small subset of IFN-responsive genes. By contrast, we observed a transcriptional profile in bystander cells that was distinct from both infected and mock cells.

In bystander cells, we observed upregulation of genes encoding cell surface receptors involved in cell-cell and extracellular matrix sensing, alongside cytoskeletal components, reminiscent of a wound healing response. Moreover, while proinflammatory cytokine expression was markedly lower than infected cells, we observed upregulation of cytokine-responsive genes, including NF-κB and AP-1. Notably, the magnitude of transcriptional changes in bystander cells was modest compared to those observed in infected cells. We therefore suggest that bystander cells exhibit a coordinated, low-amplitude response to the local inflammatory environment and tissue perturbation associated with infection. Our engineered reporter virus allows for sensitive sorting of infected and bystander cells and, consequently, the detection of subtle transcriptional differences.

Together, our tools address a critical gap in the experimental tractability of HCoV-OC43, providing a foundation for more detailed molecular and cellular studies of a historically understudied human coronavirus. These open new avenues of research in HCoV-OC43, a relevant human pathogen and a model biosafety level 2 betacoronavirus, and set a foundation for the development of new tools for other equally understudied coronaviruses.

## Acknowledgments

We acknowledge the Francis Crick Institute Flow Cytometry Science Technology Platform (STP), and particularly Sina Namjou, for their assistance with cell sorting experiments, and the Francis Crick Institute Cell Sciences STP, for supporting cell line maintenance and testing. We acknowledge the Francis Crick Institute Genomics STP, and particularly Deb Jackson, Ashley Fowler and Marg Crawford, for their contributions to mRNA library preparation and sequencing. We thank Andrew Davidson for comments and suggestions on the manuscript.

## Funding information

This work was supported by the Francis Crick Institute which receives its core funding from Cancer Research UK (CC2166, CC2277), the UK Medical Research Council (CC2166, CC2277), and the Wellcome Trust (CC2166, CC2277). This work was also supported by the UK Medical Research Council (to DLVB; MR/W005611/1, MR/Y004205/1), a University of Bristol School of Cellular and Molecular Medicine Lectureship (to HVM) and an EMBO Postdoctoral Fellowship (to NBS; ALTF 257-2022). The funders had no role in the study design, data collection and analysis, decision to publish, or preparation of the manuscript.

## Conflicts of interest

The authors declare that there are no conflicts of interest.

## Materials and Methods

### Cell culture and viruses

Cells were maintained in Dulbecco’s Modified Eagle Medium (DMEM), supplemented with 10% foetal calf serum and 100 U/mL penicillin-streptomycin, and routinely tested for Mycoplasma by The Francis Crick Institute Cell Sciences facility. Mv.1.Lu cells were a gift from Jonathan Stoye. Vero E6 (Pasteur) cells were a gift from Suzannah Rihn. MRC-5 cells were purchased from the UKHSA culture collection (05072101). Huh-7.5 (92) and Huh-7.5.1 cells (93) were generously provided by Dr Charles M Rice. HEK293T, A549 and HRT-18 cells were provided by the Francis Crick Institute Cell Sciences facility. The HCoV-OC43 isolate was purchased from ATCC (Betacoronavirus 1 VR-1558).

### Infections

Infections were carried out in DMEM supplemented with 2% foetal calf serum and all incubations were carried out at 33 °C. To generate virus stocks, 50-70% confluent Mv.1.Lu cells were inoculated with HCoV-OC43 at a multiplicity of 0.0001 PFU/cell. Cells were incubated for 3-5 days until cytopathic effect was visible and cells were starting to detach from the monolayer. Flasks were frozen at −80 °C, then thawed (94). Supernatants were clarified by centrifugation at >3,000 x g for 10 minutes, before aliquoting and storage at −80 °C.

Stocks were titrated by plaque assay on Mv.1.Lu cells. The protocol was adapted from Bracci *et al*. (56), with modifications: briefly, confluent Mv.1.Lu cells in 12-well plates were infected with serial dilutions of HCoV-OC43 in an inoculum volume of 0.25 mL for 60 minutes. Cells were then overlaid with 1 mL of overlay media: MEM (1 X), foetal calf serum (10 %), penicillin-streptomycin (1 %), Avicel (0.94 %). Dilution of overlay with inoculum results in a final Avicel concentration of 0.75%.

For assessing the stability of fluorescent reporter viruses, Mv.1.Lu cells were infected at a multiplicity of 0.0001 PFU/cell at passage 1. Subsequently, 10 µL of supernatant was passaged blind onto fresh cells every 3-5 days until cytopathic effect was observed, up to passage 5. For all other experiments, multiplicities of infection and time points for harvest are indicated in the Figures and their respective legends.

Supernatants were harvested and analysed by plaque assay on Mv.1.Lu cells. For RNA extraction, cells were lysed in TRIzol, or supernatants were mixed 1:3 with TRIzol LS (Invitrogen). RNA was extracted using the Direct-zol RNA miniprep kit, Direct-zol-96 MagBead RNA kit (Zymo Research), or by phase separation followed by isopropanol precipitation, according to the manufacturer’s protocol. For immunoblotting, cells were harvested in passive lysis buffer (Promega) supplemented with 1% IGEPAL CA-630 (Sigma-Aldrich).

To concentrate viral supernatants, 6 × 10^7^ Mv.1.Lu cells were infected at an MOI of 0.0001. Supernatants were harvested after 5 days and clarified, overlaid on top of 30% sucrose in 10 mM HEPES pH 7.5, 0.9% NaCl, and centrifuged at 100,000 x g for two hours to pellet virions. Virions were resuspended in HEPES-saline and lysed with passive lysis buffer with 1% IGEPAL CA-630 for immunoblotting.

### RT-qPCR and RT-PCR

RNA was reverse transcribed using the SuperScript VILO cDNA synthesis kit (Invitrogen) or M-MLV reverse transcriptase (Promega) using random hexamer primers.

Primer and probe sequences for HCoV-OC43 M gene have been described (57). Primers and a probe targeting nsp12 were designed to match the melting conditions and product size of the M primer-probe set, to facilitate multiplexing (Table 1). Specificity and linear signal amplification were verified using cDNA standard curves, with PCR amplicons derived from HCoV-OC43 cDNA. For growth curves, data were normalised to cellular 18S rRNA, measured using the Eukaryotic 18S rRNA Endogenous Control (VIC™/MGB probe, primer limited) (Applied Biosystems), and expressed as 2^-ΔCq^.

**Table 1.**
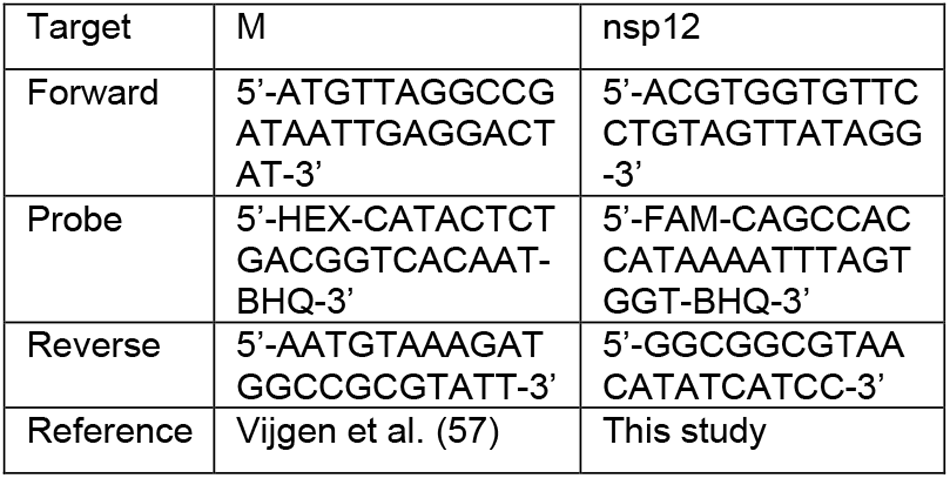
Primer and probe sequences for RT-qPCR analysis of HCoV-OC43.

For assessing the genetic stability of fluorescent tags (Figure 3H, S5D), endpoint PCR was carried out on cDNA from cell supernatants, using Platinum SuperFi PCR Master Mix (Invitrogen) using primers spanning the fluorescent insert (ORF4.5-mNG) or within the mNG coding region (dNS2-mNG). PCR cycling conditions were 95 °C for 2 min, followed by 40 cycles of 95 °C for 30 s, 60 °C for 30 s, and 72 °C for 60 s, with a final extension of 72 °C for 5 min. PCR products were separated in 1 % agarose-TBE gels and visualised using a UVP GelSolo (Analytik Jena).

### Immunoblotting

Cell lysates (5-10 ug total protein) were separated by SDS-PAGE using Any kD precast gels (Bio-Rad) and transferred to 0.45-um nitrocellulose membrane by semidry electrotransfer. Membranes were probed with sheep polyclonal anti-Nucleocapsid anti-serum (MRC PPU & CVR Coronavirus Toolkit, Sheep No. DA116, 1:1000) (48), rabbit anti-Nucleocapsid (40643-T62, Sino Biological, 1:1000), anti-Spike (PAB21478-100, The Native Antigen Company, 1:1000), anti-SARS-CoV Membrane (ABIN1887462, Antibodies Online, 1:200), anti-MHV nucleocapsid J3.3 (59) (1:200), anti-GAPDH (AM4300, Invitrogen, 1:4000) and anti-HSP90 (MA5-35624, Invitrogen, 1:2000). IR-dye labelled secondary antibodies (Li-Cor) were visualised using an Odyssey CLx imaging system (Li-Cor).

### Microscopy

To image cytopathic effect (Figure 1C, S2) cells were imaged using the ZOE Fluorescent Cell Imaging System (Bio-Rad). Fluorescent viruses (Figure 3D, S5) were imaged using an EVOS M5000 Imaging System (Invitrogen).

A549 cells grown on coverslips were infected with HCoV-OC43 or mock infected and fixed at 20 hpi in 4% formaldehyde (Sigma) for 20 min. Cells were permeabilised with 0.1% Triton X-100 (Sigma) for 3 min and blocked in 3% BSA (Sigma, A9647) for 30 min. Primary antibodies are listed under Immunoblotting above; additionally mouse anti dsRNA J2 (absolute antibodies, AB01299-2.0), mouse anti N clone 542-7D (Merck, MAB9013) and mouse anti S clone 29 (Sino Biological, 68086-MM29) were used. All primary antibodies were diluted 1: 1000 and were applied for 1 h at RT, followed by donkey anti-rabbit or anti-mouse Alexa Fluor Plus 555 or 647 secondary antibodies (Thermo Fisher) for 45 min. Nuclei were stained with Hoechst 33342 (Sigma). Coverslips were mounted in ProLong Glass (Thermo Fisher) and imaged on a VTI-SIM microscope (VisiTech). Images were processed in Fiji/ImageJ (95, 96)

### Plasmids

An archiving vector, pCLVR, was designed based on a pUC vector background, with the addition of bidirectional E. coli transcriptional terminators flanking the cloning site, to minimise read-in or read-out transcription from inserts. The cloning site consists of 5′ and 3′ BsaI and BsmBI sites, which facilitate scarless excision of inserts. The pCLVR vector contains an mRFP1_Magenta chromoprotein (97) between the cloning sites, which is excised upon fragment insertion, enabling colour-based screening of bacterial colonies.

Primers were designed across the HCoV-OC43 genome, using the ATCC reference strain, split into 11 fragments which overlap by 70 nt to allow genome assembly. Sequences were amplified from cDNA derived from HCoV-OC43-infected cells and assembled individually or in pairs into the pCLVR archive plasmid. Finally, a linker plasmid was synthesised (Genscript), containing a 30-nucleotide poly-A sequence, followed by a hepatitis delta virus ribozyme, the SV40 terminator and a PolII pause site; the linker contains a modified CMV promoter sequence, with a focussed initiator consensus sequence (98) to drive specific initiation at the 5′ end of the HCoV-OC43 genome. These sequences are flanked by ∼75 nt of the HCoV 3′ and 5′ UTRs, respectively, to facilitate assembly into a single circular plasmid. Finally, fluorescent reporters were codon optimised to match the low GC content of coronavirus genomes, and were synthesised and cloned into the pCLVR archive vector (Genscript).

### HCoV-OC43 assembly and rescue

For assembly, fragments were PCR amplified from pCLVR archive plasmid using Platinum SuperFi PCR Master Mix (Invitrogen). PCR products were purified using the NucleoSpin Gel & PCR Clean-up Kit (Macherey-Nagel) and equimolar amounts (typically 200 ng DNA per 6 kb fragment with 50 ng linker) were assembled using the NEBuilder HiFi DNA Assembly Master Mix (New England Biolabs).

Assembly reactions were stored or transfected directly into HEK293T cells, without further purification, using Lipofectamine 3000 (Invitrogen). After incubation overnight, media was exchanged for fresh DMEM supplemented with 2% foetal calf serum. Cells were then incubated for a further 3 to 7 days at 33 °C. Passage 0 supernatants were amplified on Mv.1.Lu cells to generate P1 viral stocks.

### Cell sorting

To isolate mNG-positive (infected) and mNG-negative (bystander) populations, 90-100% confluent A549 cells were inoculated with either HCoV-OC43-WT, HCoV-OC43-dNS2-mNG or HCoV-ORF4.5-mNG at an MOI of 1 PFU/cell. After incubation for 24 hours at 33 °C, cells were trypsinised, centrifuged and incubated in Zombie NIR ™ viability dye (Sony Biotechnology, 1:1000).

Cells were sorted using a BD FACSAria III cell sorter (BD Biosciences). The thresholds for detection of mNG-positive cells were set using a mock-infected control. Cell populations were sorted into collection tubes containing 2% foetal calf serum and 0.5 µl RNasin (Promega). Flow cytometry data were analysed in FlowJo software (v10.10).

### RNA sequencing

For both bulk RNA-seq and sorted RNA-seq, RNA extractions were performed by lysing cells in TRIzol (Invitrogen). RNA was extracted using the Direct-zol RNA miniprep kit or Direct-zol RNA microprep kit (Zymo Research). The Genomics Science Technology Platform at the Francis Crick Institute prepared reverse stranded sequencing libraries from 40 ng of RNA using the Watchmaker mRNA Library Prep Kit (modules K0105 and K0078), following the manufacturer’s protocols. Paired-end sequencing (100 bp) was performed on an Illumina NovaSeq X platform, with an average depth of 25 million reads per sample.

### Data analysis

Paired-end bulk RNA sequencing reads were processed using the nfcore/rnaseq pipeline (v3.10.1). The pipeline utilized STAR (99) for read alignment to the human reference genome (GRCh38.95 from Ensembl release 95, with corresponding GTF annotation v95), quantification was performed using RSEM (100). Read count matrices from the nfcore pipeline were imported into R for differential expression analysis using DESeq2 (101). The DESeq2 workflow included: filtering of low-abundance genes to improve statistical power and reduce multiple testing burden, estimation of size factors for normalization across samples, dispersion estimation and shrinkage for improved estimates of fold-changes, multiple hypothesis testing correction using the Benjamini-Hochberg false discovery rate (FDR) method.

Gene set enrichment analysis (GSEA) was performed using clusterProfiler (v4.14.6) (102) using Fast GSEA v1.32.4 (103) against curated Biological Pathway and Molecular Function databases. A 60% simplify threshold was applied to collapse redundant terms. Analyses were conducted with organism-specific annotation databases (org.Hs.eg.db for human samples). Heatmaps were visualised using pheatmap (v1.0.13 https://github.com/raivokolde/pheatmap). Gene sets for heatmaps were based on GSEA terms enriched in HCoV-OC43-ORF4.5 infected or bystander populations.

### sgRNA analysis

A subset of viral-mapping reads was further processed through a pipeline to identify discontinuous viral transcripts, including canonical subgenomic RNAs. The analysis was conceptually analogous to detecting spliced eukaryotic transcripts. Raw paired-end sequencing reads were processed using TrimGalore! (v0.6.10) to remove adapters and low-quality sequences, and reads with a minimum length of 30 bp were retained. Overlapping R1 and R2 reads were merged using FLASH (v1.2.11) (104) with a minimum overlap of 18 bp and a maximum overlap difference of 0.25. Merged reads were combined with unmerged R1 reads, which remained on the forward (R1) strand. For unmerged R2 reads, reverse complementation was performed using the FASTX Toolkit (v0.0.14, http://hannonlab.cshl.edu/fastx_toolkit/) before merging. Finally, all merged sequences (which aligned to the original R1 negative strand) were reverse complemented to the positive strand orientation for viral genome alignment.

For each viral variant, a joint reference genome was constructed by concatenating the human reference genome (GRCh38.95 from Ensembl release 95) with the corresponding viral genome sequence and annotations, and STAR indices (v2.7.11a) generated (99). Separate STAR indices were also constructed for each viral genome alone; the --genomeSAindexNbases parameter was set to 6 for viral-only indices due to the smaller genome size.

Preprocessed reads were aligned to each joint host-viral reference genome using STAR and reads mapping to viral chromosomes were extracted. Reads mapping to viral sequences were re-aligned to viral-only genomes using STAR with stringent parameters optimised for detecting chimeric reads (mapping uniquely to two regions of the genome, with a single breakpoint), corresponding to discontinuous coronavirus transcripts. Key parameters included: --outFilterType BySJout, --alignSJoverhangMin 20, --outSJfilterOverhangMin 20 20 outSJfilterCountUniqueMin 1 1 20 20, -- 1 1, -- outSJfilterCountTotalMin 1 1 1 1, -- outSJfilterDistToOtherSJmin 0 0 0 0, --scoreGapNoncan 0, --scoreGapGCAG 0, --scoreGapATAC 0, -- alignSJstitchMismatchNmax 0 0 0 0, bases -- outFilterMatchNmin 40, --outFilterMismatchNoverLmax 0.1, --alignEndsType Local, --outSAMmultNmax 1, -- alignSJDBoverhangMin 20.

BAM files from viral-specific alignments were processed using GenomicAlignments in R (105) to extract alignment information. The number of chimeric and nonchimeric alignments were counted for each sample. A background measure of nonchimeric (i.e. genomic) reads was defined for each sample to account for variation in viral RNA abundance across samples. The genomic background region was defined as a 1kb window centred on the midpoint between the start coordinate of the S sub-genomic RNA (sgRNA) intron and the start of the ORF1ab gene body. Nonchimeric reads that overlapped this region were counted as “genomic” alignments and used as a normalization factor for subsequent analyses. Transcript junctions were extracted from chimeric alignments using the ‵summarizeJunctions()′ function, which identified all unique junction coordinates and their read depths.

Junctions were classified as either canonical (donor and acceptor sites match coordinates in the annotated sgRNA GTF), alternative (only one donor or acceptor site match), or defective (neither donor nor acceptor site match). Within the canonical class, junctions were further subclassified by their corresponding sgRNA gene identity (e.g., E, M, N). Defective junctions were subclassified based on the size of their genomic span (gap size).

### Phylogenetic tree

The ORF1ab sequence was extracted from representative coronavirus genome sequences (NC_005831.2, NC_039208.1, NC_004718.3, NC_006213.1, NC_002645.1, NC_048217.1, NC_045512.2, NC_006577.2, NC_019843.3, NC_048213.1, NC_038861.1, NC_002306.3). Protein alignments were made using MAFFT (106) and trees were constructed using FastTree (107), visualised using ape (doi:10.1093/bioinformatics/bty633) and ggtree (108).

## Notes

### Competing Interest Statement

The authors have declared no competing interest.

## References

1. Holmes EC. The Emergence and Evolution of SARS-CoV-2. Annu Rev Virol. 2024;11(1):21–42.

2. de Wit E, van Doremalen N, Falzarano D, Munster VJ. SARS and MERS: recent insights into emerging coronaviruses. Nat Rev Microbiol. 2016;14(8):523–34.

3. Buonavoglia A, Pellegrini F, Decaro N, Galgano M, Pratelli A. A One Health Perspective on Canine Coronavirus: A Wolf in Sheep’s Clothing? Microorganisms. 2023;11(4).

4. Zehr JD, Pond SLK, Martin DP, Ceres K, Whittaker GR, Millet JK, et al. Recent Zoonotic Spillover and Tropism Shift of a Canine Coronavirus Is Associated with Relaxed Selection and Putative Loss of Function in NTD Subdomain of Spike Protein. Viruses. 2022;14(5).

5. Lau SK, Lee P, Tsang AK, Yip CC, Tse H, Lee RA, et al. Molecular epidemiology of human coronavirus OC43 reveals evolution of different genotypes over time and recent emergence of a novel genotype due to natural recombination. J Virol. 2011;85(21):11325–37.

6. Wilson R, Kovacs D, Crosby M, Ho A. Global Epidemiology and Seasonality of Human Seasonal Coronaviruses: A Systematic Review. Open Forum Infect Dis. 2024;11(8):ofae418.

7. Kesheh MM, Hosseini P, Soltani S, Zandi M. An overview on the seven pathogenic human coronaviruses. Rev Med Virol. 2022;32(2):e2282.

8. Dijkman R, Jebbink MF, Gaunt E, Rossen JW, Templeton KE, Kuijpers TW, et al. The dominance of human coronavirus OC43 and NL63 infections in infants. J Clin Virol. 2012;53(2):135–9.

9. Zeng ZQ, Chen DH, Tan WP, Qiu SY, Xu D, Liang HX, et al. Epidemiology and clinical characteristics of human coronaviruses OC43, 229E, NL63, and HKU1: a study of hospitalized children with acute respiratory tract infection in Guangzhou, China. Eur J Clin Microbiol Infect Dis. 2018;37(2):363–9.

10. Falsey AR, McCann RM, Hall WJ, Criddle MM, Formica MA, Wycoff D, et al. The “common cold” in frail older persons: impact of rhinovirus and coronavirus in a senior daycare center. J Am Geriatr Soc. 1997;45(6):706–11.

11. Ohyama K, Honda H, Aoki M, Wakuda M, Kitahara T, Kaede C, et al. Resurgence of human coronavirus OC43 at a long-term care facility during the coronavirus disease 2019 (COVID-19) pandemic: Outbreak investigation. Antimicrob Steward Healthc Epidemiol. 2023;3(1):e97.

12. Lau SKP, Li KSM, Li X, Tsang KY, Sridhar S, Woo PCY. Fatal Pneumonia Associated With a Novel Genotype of Human Coronavirus OC43. Front Microbiol. 2021;12:795449.

13. Kasereka MC, Hawkes MT. Neuroinvasive potential of human coronavirus OC43: case report of fatal encephalitis in an immunocompromised host. J Neurovirol. 2021;27(2):340–4.

14. St-Jean JR, Jacomy H, Desforges M, Vabret A, Freymuth F, Talbot PJ. Human respiratory coronavirus OC43: genetic stability and neuroinvasion. J Virol. 2004;78(16):8824–34.

15. Arbour N, Day R, Newcombe J, Talbot PJ. Neuroinvasion by human respiratory coronaviruses. J Virol. 2000;74(19):8913–21.

16. St-Jean JR, Desforges M, Almazan F, Jacomy H, Enjuanes L, Talbot PJ. Recovery of a neurovirulent human coronavirus OC43 from an infectious cDNA clone. J Virol. 2006;80(7):3670–4.

17. Pyrc K, Sims AC, Dijkman R, Jebbink M, Long C, Deming D, et al. Culturing the unculturable: human coronavirus HKU1 infects, replicates, and produces progeny virions in human ciliated airway epithelial cell cultures. J Virol. 2010;84(21):11255–63.

18. Dijkman R, Jebbink MF, Koekkoek SM, Deijs M, Jonsdottir HR, Molenkamp R, et al. Isolation and characterization of current human coronavirus strains in primary human epithelial cell cultures reveal differences in target cell tropism. J Virol. 2013;87(11):6081–90.

19. Schirtzinger EE, Kim Y, Davis AS. Improving human coronavirus OC43 (HCoV-OC43) research comparability in studies using HCoV-OC43 as a surrogate for SARS-CoV-2. J Virol Methods. 2022;299:114317.

20. Fausto A, Otter CJ, Bracci N, Weiss SR. Improved Culture Methods for Human Coronaviruses HCoV-OC43, HCoV-229E, and HCoV-NL63. Curr Protoc. 2023;3(10):e914.

21. Hirose R, Watanabe N, Bandou R, Yoshida T, Daidoji T, Naito Y, et al. A Cytopathic Effect-Based Tissue Culture Method for HCoV-OC43 Titration Using TMPRSS2-Expressing VeroE6 Cells. mSphere. 2021;6(3).

22. Almazan F, Sola I, Zuniga S, Marquez-Jurado S, Morales L, Becares M, et al. Coronavirus reverse genetic systems: infectious clones and replicons. Virus Res. 2014;189:262–70.

23. Masters PS. Reverse genetics of the largest RNA viruses. Adv Virus Res. 1999;53:245–64.

24. Thiel V, Siddell SG. Reverse genetics of coronaviruses using vaccinia virus vectors. Curr Top Microbiol Immunol. 2005;287:199–227.

25. Masters PS, Rottier PJ. Coronavirus reverse genetics by targeted RNA recombination. Curr Top Microbiol Immunol. 2005;287:133–59.

26. Yount B, Curtis KM, Baric RS. Strategy for systematic assembly of large RNA and DNA genomes: transmissible gastroenteritis virus model. J Virol. 2000;74(22):10600–11.

27. Yount B, Denison MR, Weiss SR, Baric RS. Systematic assembly of a full-length infectious cDNA of mouse hepatitis virus strain A59. J Virol. 2002;76(21):11065–78.

28. Almazan F, Gonzalez JM, Penzes Z, Izeta A, Calvo E, Plana-Duran J, et al. Engineering the largest RNA virus genome as an infectious bacterial artificial chromosome. Proc Natl Acad Sci U S A. 2000;97(10):5516–21.

29. Thi Nhu Thao T, Labroussaa F, Ebert N, V’Kovski P, Stalder H, Portmann J, et al. Rapid reconstruction of SARS-CoV-2 using a synthetic genomics platform. Nature. 2020;582(7813):561–5.

30. Duguay BA, Tooley TH, Pringle ES, Rohde JR, McCormick C. A yeast-based reverse genetics system to generate HCoV-OC43 reporter viruses encoding an eighth subgenomic RNA. J Virol. 2025;99(2):e0167124.

31. Li Y, Duan L, Tang L, Huang M, Zhao Y, Zhang G, et al. Rapid reconstruction of infectious bronchitis virus expressing fluorescent protein from its nsp2 gene based on transformation-associated recombination platform. J Virol. 2025;99(7):e0053525.

32. Ye F, Wang N, Guan Q, Wang M, Sun J, Zhai D, et al. Rapid generation and characterization of recombinant HCoV-OC43-VR1558 infectious clones expressing reporter Renilla luciferase. Biosaf Health. 2024;6(6):350–60.

33. Ujike M, Etoh Y, Urushiyama N, Taguchi F, Asanuma H, Enjuanes L, et al. Reverse Genetics with a Full-Length Infectious cDNA Clone of Bovine Torovirus. J Virol. 2022;96(3):e0156121.

34. Thiel V, Herold J, Schelle B, Siddell SG. Infectious RNA transcribed in vitro from a cDNA copy of the human coronavirus genome cloned in vaccinia virus. J Gen Virol. 2001;82(Pt 6):1273–81.

35. Coley SE, Lavi E, Sawicki SG, Fu L, Schelle B, Karl N, et al. Recombinant mouse hepatitis virus strain A59 from cloned, full-length cDNA replicates to high titers in vitro and is fully pathogenic in vivo. J Virol. 2005;79(5):3097–106.

36. van den Worm SH, Eriksson KK, Zevenhoven JC, Weber F, Zust R, Kuri T, et al. Reverse genetics of SARS-related coronavirus using vaccinia virus-based recombination. PLoS One. 2012;7(3):e32857.

37. Kurhade C, Xie X, Shi PY. Reverse genetic systems of SARS-CoV-2 for antiviral research. Antiviral Res. 2023;210:105486.

38. Torii S, Ono C, Suzuki R, Morioka Y, Anzai I, Fauzyah Y, et al. 2020.

39. Melade J, Piorkowski G, Touret F, Fourie T, Driouich JS, Cochin M, et al. A simple reverse genetics method to generate recombinant coronaviruses. EMBO Rep. 2022;23(5):e53820.

40. Ujike M, Suzuki T. Progress of research on coronaviruses and toroviruses in large domestic animals using reverse genetics systems. Vet J. 2024;305:106122.

41. Jiang H, Wang T, Kong L, Li B, Peng Q. Reverse Genetics Systems for Emerging and Re-Emerging Swine Coronaviruses and Applications. Viruses. 2023;15(10).

42. Bilotti K, Keep S, Sikkema AP, Pryor JM, Kirk J, Foldes K, et al. One-pot Golden Gate Assembly of an avian infectious bronchitis virus reverse genetics system. PLoS One. 2024;19(7):e0307655.

43. Torii S, Ono C, Suzuki R, Morioka Y, Anzai I, Fauzyah Y, et al. Establishment of a reverse genetics system for SARS-CoV-2 using circular polymerase extension reaction. Cell Rep. 2021;35(3):109014.

44. Freeman MC, Graham RL, Lu X, Peek CT, Denison MR. Coronavirus replicase-reporter fusions provide quantitative analysis of replication and replication complex formation. J Virol. 2014;88(10):5319–27.

45. V’Kovski P, Gerber M, Kelly J, Pfaender S, Ebert N, Braga Lagache S, et al. Determination of host proteins composing the microenvironment of coronavirus replicase complexes by proximity-labeling. Elife. 2019;8.

46. de Haan CA, Haijema BJ, Boss D, Heuts FW, Rottier PJ. Coronaviruses as vectors: stability of foreign gene expression. J Virol. 2005;79(20):12742–51.

47. Bentley K, Armesto M, Britton P. Infectious Bronchitis Virus as a Vector for the Expression of Heterologous Genes. PLoS One. 2013;8(6):e67875.

48. Rihn SJ, Merits A, Bakshi S, Turnbull ML, Wickenhagen A, Alexander AJT, et al. A plasmid DNA-launched SARS-CoV-2 reverse genetics system and coronavirus toolkit for COVID-19 research. PLoS Biol. 2021;19(2):e3001091.

49. Khan JQ, Rohamare M, Rajamanickam K, Bhanumathy KK, Lew J, Kumar A, et al. Generation of a SARS-CoV-2 Reverse Genetics System and Novel Human Lung Cell Lines That Exhibit High Virus-Induced Cytopathology. Viruses. 2023;15(6).

50. Hou YJ, Okuda K, Edwards CE, Martinez DR, Asakura T, Dinnon KH, 3rd, et al. SARS-CoV-2 Reverse Genetics Reveals a Variable Infection Gradient in the Respiratory Tract. Cell. 2020;182(2):429–46 e14.

51. Sola I, Almazan F, Zuniga S, Enjuanes L. Continuous and Discontinuous RNA Synthesis in Coronaviruses. Annu Rev Virol. 2015;2(1):265–88.

52. Fischer F, Stegen CF, Koetzner CA, Masters PS. Analysis of a recombinant mouse hepatitis virus expressing a foreign gene reveals a novel aspect of coronavirus transcription. J Virol. 1997;71(7):5148–60.

53. Pasternak AO, Spaan WJ, Snijder EJ. Regulation of relative abundance of arterivirus subgenomic mRNAs. J Virol. 2004;78(15):8102–13.

54. Hsue B, Masters PS. Insertion of a new transcriptional unit into the genome of mouse hepatitis virus. J Virol. 1999;73(7):6128–35.

55. Alonso S, Izeta A, Sola I, Enjuanes L. Transcription regulatory sequences and mRNA expression levels in the coronavirus transmissible gastroenteritis virus. J Virol. 2002;76(3):1293–308.

56. Bracci N, Pan HC, Lehman C, Kehn-Hall K, Lin SC. Improved plaque assay for human coronaviruses 229E and OC43. PeerJ. 2020;8:e10639.

57. Vijgen L, Keyaerts E, Moes E, Maes P, Duson G, Van Ranst M. Development of one-step, real-time, quantitative reverse transcriptase PCR assays for absolute quantitation of human coronaviruses OC43 and 229E. J Clin Microbiol. 2005;43(11):5452–6.

58. Wurm T, Chen H, Hodgson T, Britton P, Brooks G, Hiscox JA. Localization to the nucleolus is a common feature of coronavirus nucleoproteins, and the protein may disrupt host cell division. J Virol. 2001;75(19):9345–56.

59. Fleming JO, Stohlman SA, Harmon RC, Lai MM, Frelinger JA, Weiner LP. Antigenic relationships of murine coronaviruses: analysis using monoclonal antibodies to JHM (MHV-4) virus. Virology. 1983;131(2):296–307.

60. Bost AG, Carnahan RH, Lu XT, Denison MR. Four proteins processed from the replicase gene polyprotein of mouse hepatitis virus colocalize in the cell periphery and adjacent to sites of virion assembly. J Virol. 2000;74(7):3379–87.

61. Klumperman J, Locker JK, Meijer A, Horzinek MC, Geuze HJ, Rottier PJ. Coronavirus M proteins accumulate in the Golgi complex beyond the site of virion budding. J Virol. 1994;68(10):6523–34.

62. Wang J, Fang S, Xiao H, Chen B, Tam JP, Liu DX. Interaction of the coronavirus infectious bronchitis virus membrane protein with beta-actin and its implication in virion assembly and budding. PLoS One. 2009;4(3):e4908.

63. Labonte P, Mounir S, Talbot PJ. Sequence and expression of the ns2 protein gene of human coronavirus OC43. J Gen Virol. 1995;76 (Pt 2):431–5.

64. Goldstein SA, Thornbrough JM, Zhang R, Jha BK, Li Y, Elliott R, et al. Lineage A Betacoronavirus NS2 Proteins and the Homologous Torovirus Berne pp1a Carboxy-Terminal Domain Are Phosphodiesterases That Antagonize Activation of RNase L. J Virol. 2017;91(5).

65. Schwarz B, Routledge E, Siddell SG. Murine coronavirus nonstructural protein ns2 is not essential for virus replication in transformed cells. J Virol. 1990;64(10):4784–91.

66. Bhargava A, Szachnowski U, Chazal M, Foretek D, Caval V, Aicher SM, et al. Transcriptomic analysis of sorted lung cells revealed a proviral activity of the NF-kappaB pathway toward SARS-CoV-2. iScience. 2023;26(12):108449.

67. Grunwell JR, Yeligar SM, Stephenson S, Ping XD, Gauthier TW, Fitzpatrick AM, et al. TGF-beta1 Suppresses the Type I IFN Response and Induces Mitochondrial Dysfunction in Alveolar Macrophages. J Immunol. 2018;200(6):2115–28.

68. Guerin MV, Regnier F, Feuillet V, Vimeux L, Weiss JM, Bismuth G, et al. TGFbeta blocks IFNalpha/beta release and tumor rejection in spontaneous mammary tumors. Nat Commun. 2019;10(1):4131.

69. Shen L, Yang Y, Ye F, Liu G, Desforges M, Talbot PJ, et al. Safe and Sensitive Antiviral Screening Platform Based on Recombinant Human Coronavirus OC43 Expressing the Luciferase Reporter Gene. Antimicrob Agents Chemother. 2016;60(9):5492–503.

70. Diefenbacher MV, Baric TJ, Martinez DR, Baric RS, Catanzaro NJ, Sheahan TP. A nano-luciferase expressing human coronavirus OC43 for countermeasure development. Virus Res. 2024;339:199286.

71. Zhang Y, Chen J, Feng H, Yang L, Hu L, Gao Y, et al. Rapid generation of HCoV-229E and HCoV-OC43 reporter viruses and replicons for antiviral research. Front Cell Infect Microbiol. 2025;15:1614369.

72. Duguay BA, McCormick C. Assembly and Mutagenesis of Human Coronavirus OC43 Genomes in Yeast via Transformation-Associated Recombination. Bio Protoc. 2025;15(16):e5422.

73. Brant AC, Hu Z, Chen AZ, Majerciak V, Yewdell J, Zheng ZM. Discontinuous template switching generates coronavirus subgenomic RNAs from the 3’ viral genome end by 5’ to 3’ transcription. J Virol. 2025;99(10):e0143825.

74. Deng X, Zou J, Liang Z, Ren P, Shi PY, Menachery VD, et al. A dual-reporter HCoV-OC43 for coronavirus biology and countermeasure development. Antiviral Res. 2025;244:106306.

75. Sawicki SG, Sawicki DL. Coronaviruses use discontinuous extension for synthesis of subgenome-length negative strands. Adv Exp Med Biol. 1995;380:499–506.

76. Makino S, Soe LH, Shieh CK, Lai MM. Discontinuous transcription generates heterogeneity at the leader fusion sites of coronavirus mRNAs. J Virol. 1988;62(10):3870–3.

77. Zuniga S, Sola I, Alonso S, Enjuanes L. Sequence motifs involved in the regulation of discontinuous coronavirus subgenomic RNA synthesis. J Virol. 2004;78(2):980–94.

78. Mateos-Gomez PA, Morales L, Zuniga S, Enjuanes L, Sola I. Long-distance RNA-RNA interactions in the coronavirus genome form high-order structures promoting discontinuous RNA synthesis during transcription. J Virol. 2013;87(1):177–86.

79. Dufour D, Mateos-Gomez PA, Enjuanes L, Gallego J, Sola I. Structure and functional relevance of a transcription-regulating sequence involved in coronavirus discontinuous RNA synthesis. J Virol. 2011;85(10):4963–73.

80. Sola I, Moreno JL, Zuniga S, Alonso S, Enjuanes L. Role of nucleotides immediately flanking the transcription-regulating sequence core in coronavirus subgenomic mRNA synthesis. J Virol. 2005;79(4):2506–16.

81. Bresson S, Sani E, Armatowska A, Dixon C, Tollervey D. The transcriptional and translational landscape of HCoV-OC43 infection. PLoS Pathog. 2025;21(1):e1012831.

82. Blanco-Melo D, Nilsson-Payant BE, Liu WC, Uhl S, Hoagland D, Moller R, et al. Imbalanced Host Response to SARS-CoV-2 Drives Development of COVID-19. Cell. 2020;181(5):1036–45 e9.

83. Roth-Cross JK, Stokes H, Chang G, Chua MM, Thiel V, Weiss SR, et al. Organ-specific attenuation of murine hepatitis virus strain A59 by replacement of catalytic residues in the putative viral cyclic phosphodiesterase ns2. J Virol. 2009;83(8):3743–53.

84. Zhao L, Jha BK, Wu A, Elliott R, Ziebuhr J, Gorbalenya AE, et al. Antagonism of the interferon-induced OAS-RNase L pathway by murine coronavirus ns2 protein is required for virus replication and liver pathology. Cell Host Microbe. 2012;11(6):607–16.

85. Sui B, Huang J, Jha BK, Yin P, Zhou M, Fu ZF, et al. Crystal structure of the mouse hepatitis virus ns2 phosphodiesterase domain that antagonizes RNase L activation. J Gen Virol. 2016;97(4):880–6.

86. Irigoyen N, Firth AE, Jones JD, Chung BY, Siddell SG, Brierley I. High-Resolution Analysis of Coronavirus Gene Expression by RNA Sequencing and Ribosome Profiling. PLoS Pathog. 2016;12(2):e1005473.

87. Echavarria-Consuegra L, Cook GM, Busnadiego I, Lefevre C, Keep S, Brown K, et al. Manipulation of the unfolded protein response: A pharmacological strategy against coronavirus infection. PLoS Pathog. 2021;17(6):e1009644.

88. Fung TS, Huang M, Liu DX. Coronavirus-induced ER stress response and its involvement in regulation of coronavirus-host interactions. Virus Res. 2014;194:110–23.

89. Yu X, Abbas-Aghababazadeh F, Chen YA, Fridley BL. Statistical and Bioinformatics Analysis of Data from Bulk and Single-Cell RNA Sequencing Experiments. Methods Mol Biol. 2021;2194:143–75.

90. Hwang B, Lee JH, Bang D. Single-cell RNA sequencing technologies and bioinformatics pipelines. Exp Mol Med. 2018;50(8):1–14.

91. Kolodziejczyk AA, Kim JK, Svensson V, Marioni JC, Teichmann SA. The technology and biology of single-cell RNA sequencing. Mol Cell. 2015;58(4):610–20.

92. Blight KJ, McKeating JA, Rice CM. Highly permissive cell lines for subgenomic and genomic hepatitis C virus RNA replication. J Virol. 2002;76(24):13001–14.

93. Zhong J, Gastaminza P, Cheng G, Kapadia S, Kato T, Burton DR, et al. Robust hepatitis C virus infection in vitro. Proc Natl Acad Sci U S A. 2005;102(26):9294–9.

94. Leibowitz J, Kaufman G, Liu P. Coronaviruses: propagation, quantification, storage, and construction of recombinant mouse hepatitis virus. Curr Protoc Microbiol. 2011;Chapter 15(1):Unit 15E 1.

95. Schindelin J, Arganda-Carreras I, Frise E, Kaynig V, Longair M, Pietzsch T, et al. Fiji: an open-source platform for biological-image analysis. Nat Methods. 2012;9(7):676–82.

96. Schneider CA, Rasband WS, Eliceiri KW. NIH Image to ImageJ: 25 years of image analysis. Nat Methods. 2012;9(7):671–5.

97. Bao L, Menon PNK, Liljeruhm J, Forster AC. Overcoming chromoprotein limitations by engineering a red fluorescent protein. Anal Biochem. 2020;611:113936.

98. Vo Ngoc L, Cassidy CJ, Huang CY, Duttke SH, Kadonaga JT. The human initiator is a distinct and abundant element that is precisely positioned in focused core promoters. Genes Dev. 2017;31(1):6–11.

99. Dobin A, Davis CA, Schlesinger F, Drenkow J, Zaleski C, Jha S, et al. STAR: ultrafast universal RNA-seq aligner. Bioinformatics. 2013;29(1):15–21.

100. Li B, Dewey CN. RSEM: accurate transcript quantification from RNA-Seq data with or without a reference genome. BMC Bioinformatics. 2011;12:323.

101. Love MI, Huber W, Anders S. Moderated estimation of fold change and dispersion for RNA-seq data with DESeq2. Genome Biol. 2014;15(12):550.

102. Yu G, Wang LG, Han Y, He QY. clusterProfiler: an R package for comparing biological themes among gene clusters. OMICS. 2012;16(5):284–7.

103. Korotkevich G, Sukhov V, Budin N, Shpak B, Artyomov MN, Sergushichev A. Fast gene set enrichment analysis. bioRxiv. 2021:060012.

104. Magoc T, Salzberg SL. FLASH: fast length adjustment of short reads to improve genome assemblies. Bioinformatics. 2011;27(21):2957–63.

105. Lawrence M, Huber W, Pages H, Aboyoun P, Carlson M, Gentleman R, et al. Software for computing and annotating genomic ranges. PLoS Comput Biol. 2013;9(8):e1003118.

106. Katoh K, Standley DM. MAFFT multiple sequence alignment software version 7: improvements in performance and usability. Mol Biol Evol. 2013;30(4):772–80.

107. Price MN, Dehal PS, Arkin AP. FastTree: computing large minimum evolution trees with profiles instead of a distance matrix. Mol Biol Evol. 2009;26(7):1641–50.

108. Yu G, Smith DK, Zhu H, Guan Y, Lam TT-Y. ggtree: an r package for visualization and annotation of phylogenetic trees with their covariates and other associated data. Methods in Ecology and Evolution. 2017;8(1):28–36.

