## Supplementary Figures for "A stable subgenomic reporter coronavirus enables transcriptional profiling of bystander cells"

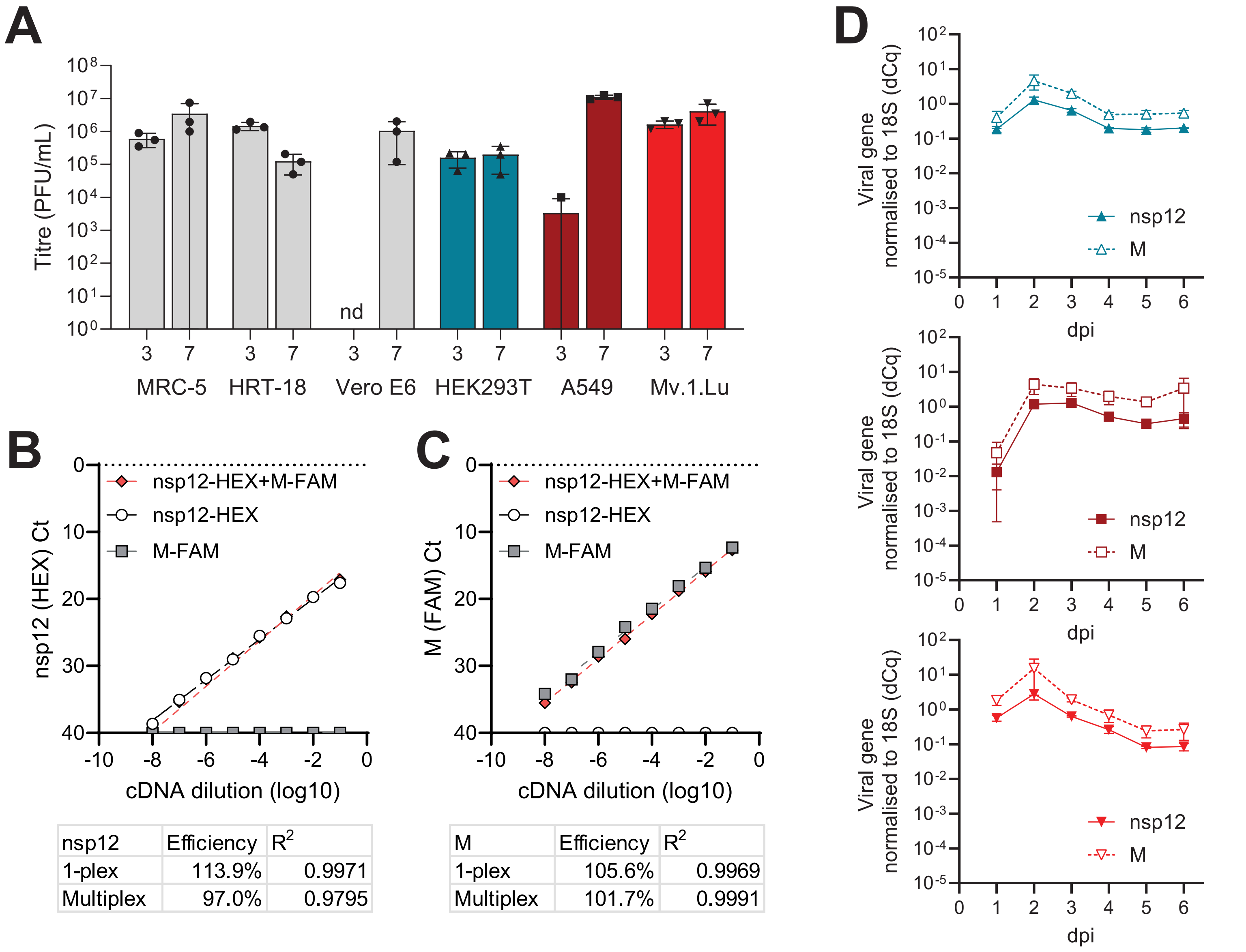


| nsp12 | Efficiency | $R^2$ |
|---|---|---|
| 1-plex | 113.9% | 0.9971 |
| Multiplex | 97.0% | 0.9795 |

| M | Efficiency | $R^2$ |
|---|---|---|
| 1-plex | 105.6% | 0.9969 |
| Multiplex | 101.7% | 0.9991 |

**Figure S1.** Growth and analysis of HCoV-OC43 in cell culture. **A.** Infectious titres of HCoV-OC43 released from the indicated cell lines at 3- and 7-days post infection (dpi) in plaque forming units (PFU)/mL. Data are means and standard deviations of three biological replicates. nd, not detected. **B-C.** Standard curves of a HEX-labelled nsp12 primer-probe set (B) and FAM-labelled M gene primer-probe set (C), either alone (white, grey) or multiplexed (red), using gene-specific PCR amplicons, amplified from cDNA from HCoV-OC43-infected cells, as templates, to test for specificity. Primer efficiency ($E = -1 + 10^{(-1/Slope)}$) and $R^2$ values are shown. **D.** Growth of HCoV-OC43 in 293T (upper), A549 (middle) or Mv.1.Lu (lower) cells infected at an MOI of 0.05, analysed by RT-qPCR. Viral gene expression (nsp12, solid line and M, dashed line) was normalised to host 18S RNA ($\Delta$Cq). dpi, days post infection. Data are means and standard deviations of at least three biological replicates.

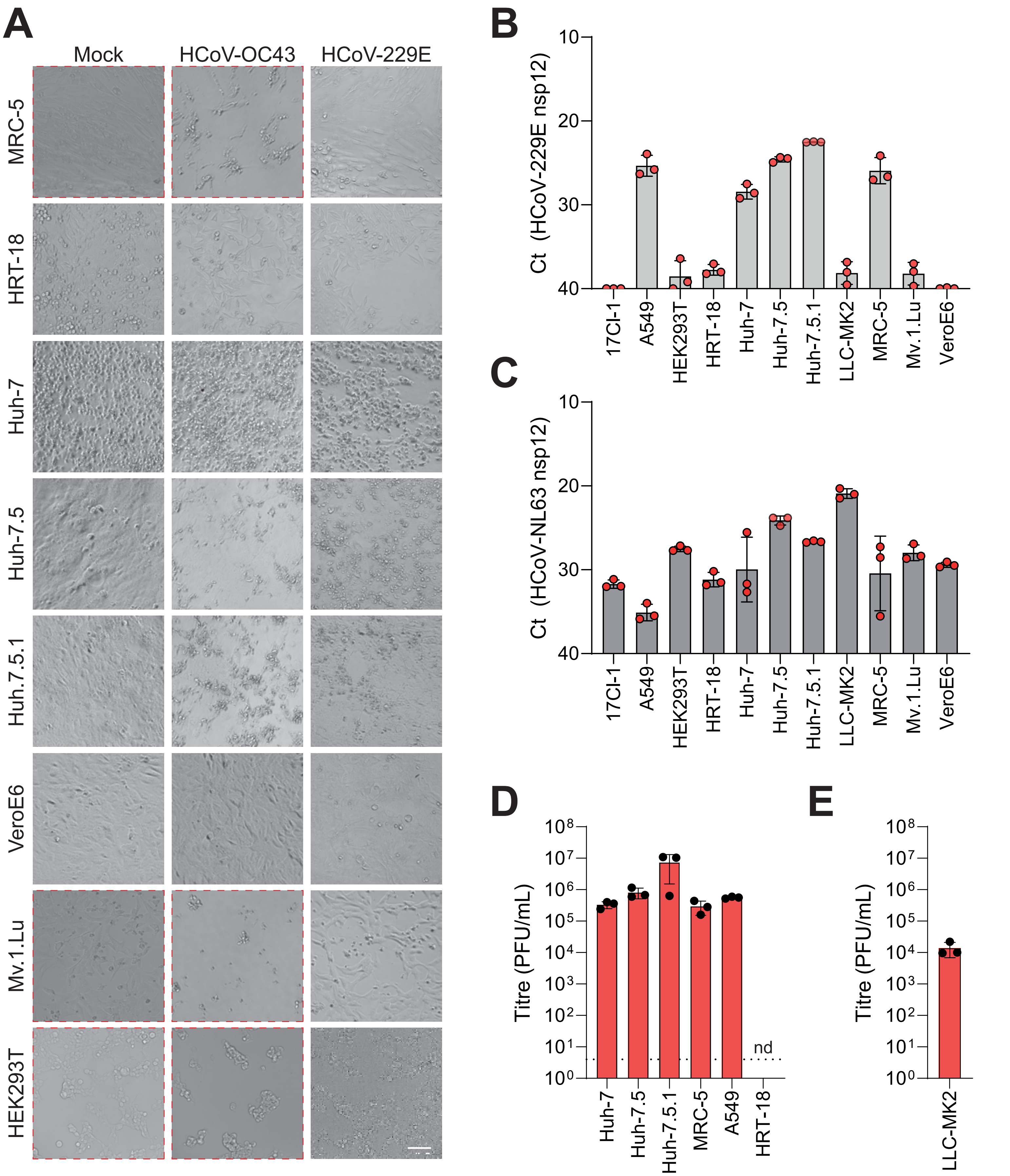


**Figure S2.** Cell culture conditions for human alphacoronaviruses. **A.** Light microscopy showing eight cell lines infected with HCoV-229E, or mock infected, at an MOI of 0.0001, five days post infection. HCoV-OC43 is included for comparison and dashed boxes indicate panels which also appear in Figure 1C and are included here for reference. Scale bar represents 100 μm. **B-C.** RT-qPCR analysis of supernatants from HCoV-229E (B) or HCoV-NL63 (C) -infected cells, five days post infection. Data represent means and standard deviations of three biological replicates. **D-E.** Plaque assay of supernatants from B and C, respectively. Plaque assays for HCoV-229E (D) were performed in Huh-7 cells and for HCoV-NL63 (E) in LLC-MK2 cells. nd, not detected.

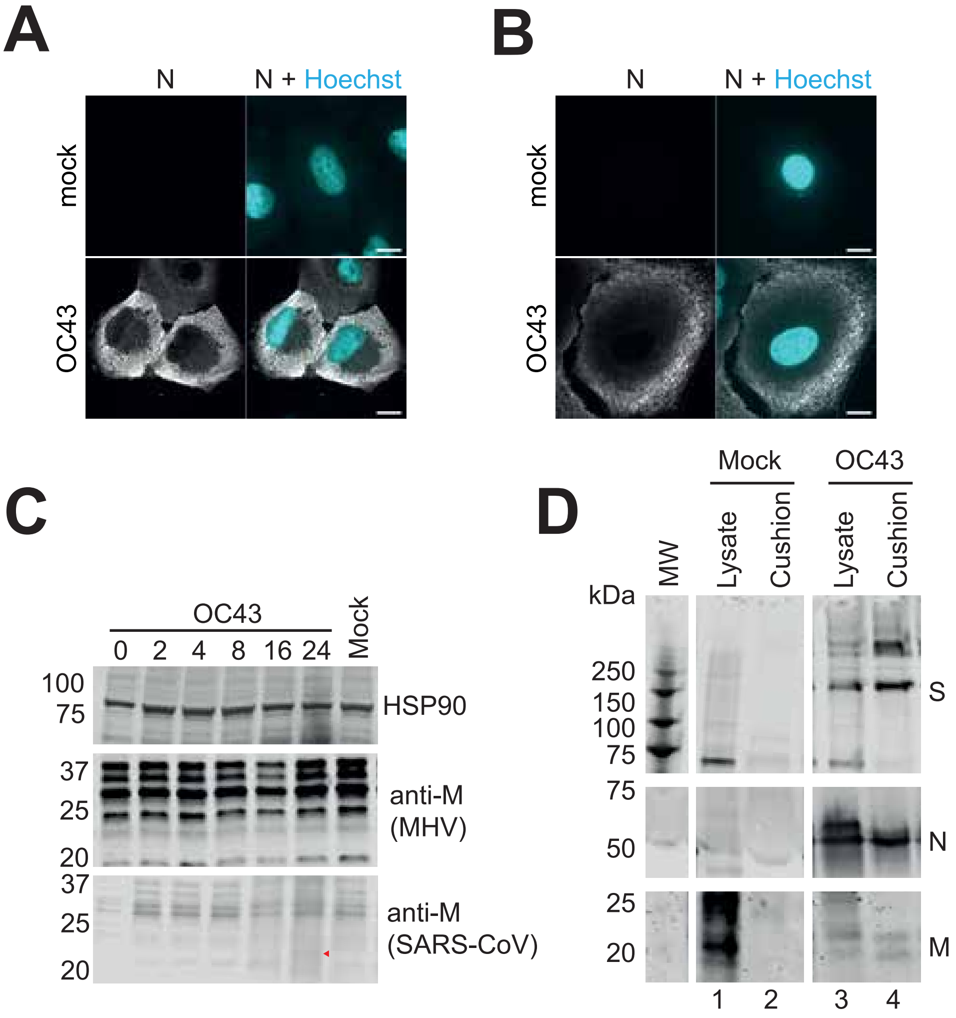


**Figure S3.** Antibody validation. **A-B.** Immunofluorescence microscopy of HCoV-OC43-infected A549 cells using two different anti-N antibodies: a rabbit polyclonal (A, SinoBiological 40643-T62, 1:1000) and a mouse monoclonal (B, Merck MAB9013, 1:1000). Scale bars represent 10 μm. **C.** Western blot analysis of Mv.1.Lu cells infected with HCoV-OC43 at an MOI of 10 for up to 24 hours, probed with two anti-M antibodies, a mouse monoclonal raised against MHV and a rabbit polyclonal raised against SARS-CoV. Weak M detection at 24 hpi is indicated by a red arrowhead. **D.** Western blot of viral supernatants concentrated by ultracentrifugation through a 30% sucrose cushion, compared to cell lysates. Background signal is higher in mock-infected cell lysates (lane 1), due to substantial cytopathic effect in infected cells.

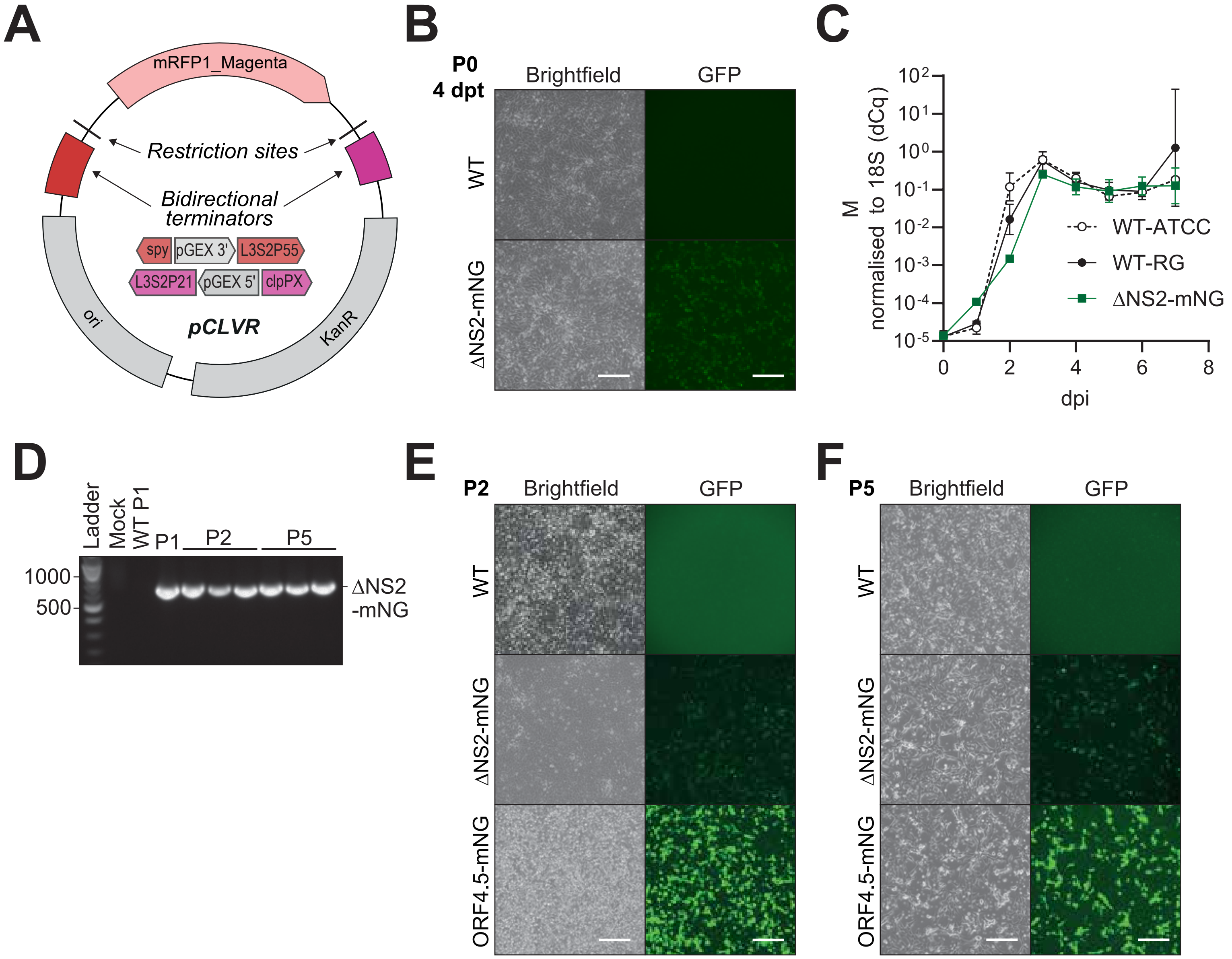


**Figure S4.** Rescue and passage of reverse genetics-derived HCoV-OC43. **A.** Schematic of the pCLVR storage vector, showing restriction sites flanking an mRFP1_Magenta chromoprotein, which is excised upon insert cloning. Upstream (red) and downstream (pink) bidirectional transcriptional terminators, including pGEX primer binding sites for plasmid sequencing, as well as bacterial replication (ori) and antibiotic resistance (KanR) elements, are indicated. **B.** Brightfield and fluorescence microscopy analysis of HEK293T cells transfected with *in vitro* assembled HCoV-OC43 infectious clones, four days post transfection. **C.** Replication of WT (ATCC isolate, dashed line) and RG-derived HCoV-OC43 in Mv.1.Lu cells (MOI 0.01) measured by RT-qPCR against the viral M gene, normalised to host 18S rRNA (ΔCq). Data are means and standard deviations of three biological replicates. **D.** Agarose gel analysis of PCR amplicons within the ΔNS2-mNG insertion site, amplified from RNA purified from supernatants, from passage 1 (P1), P2 or P5. Three biological replicates of P2 and P5 were analysed. **E-F.** Brightfield and fluorescence microscopy analysis of Mv.1.Lu cells infected with WT and fluorescent reporter viruses at passage 2 (P2, C) and P5 (D). Scale bars represent 200 μm.

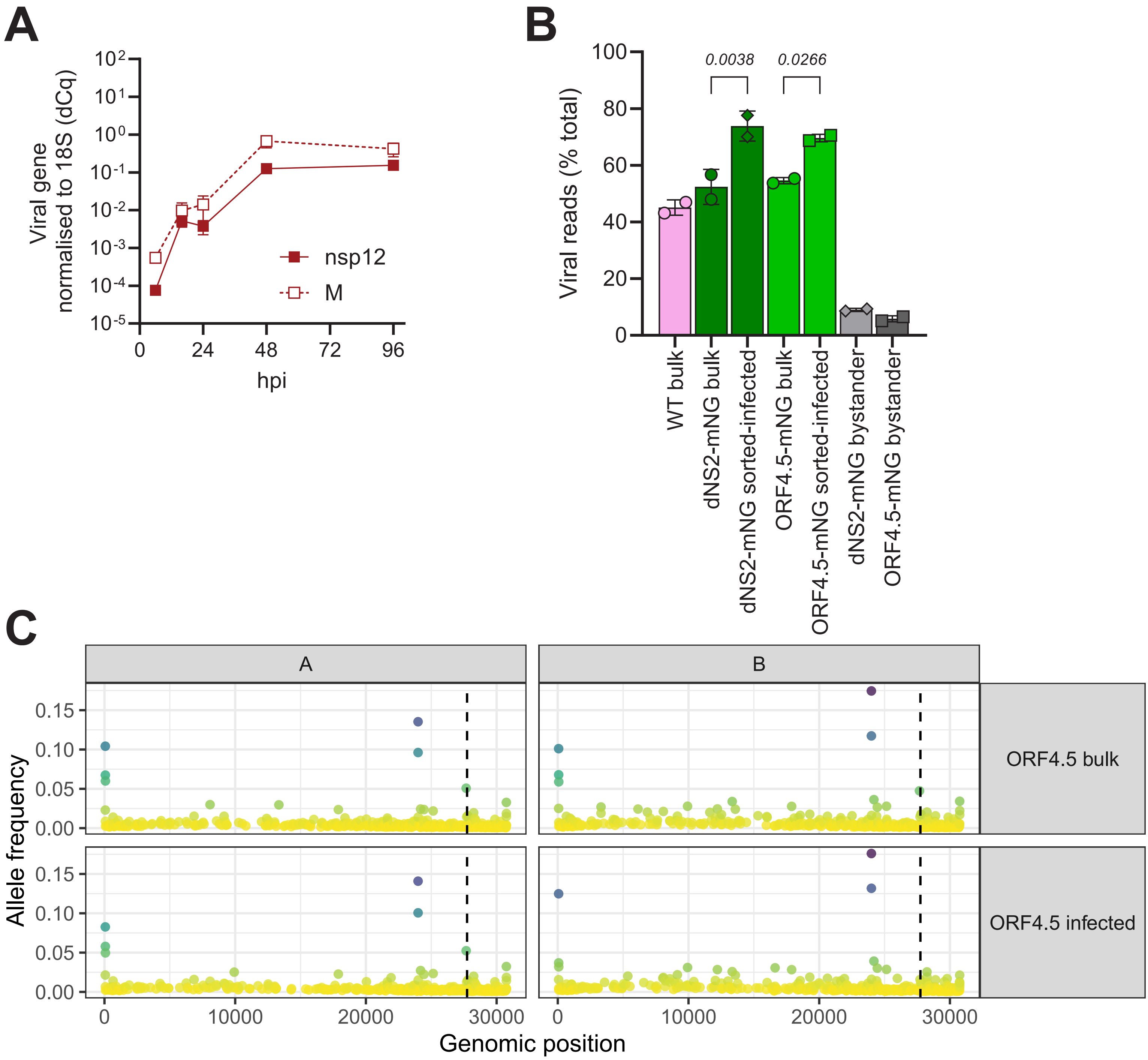


**Figure S5.** Transcriptomic analysis of HCoV-OC43-infected A549 cells. **A.** Growth of HCoV-OC43 in A549 cells (MOI 1), analysed by RT-qPCR. Viral gene expression (nsp12, solid line and M, dashed line) was normalised to host 18S RNA ($\Delta$Cq). dpi, days post infection. Data are means and standard deviations of three biological replicates. **B.** Proportion of viral reads in infected A549 cells, from bulk and sorted cell populations. Data represent means and standard deviations of two independent experiments. Means were compared by one-way ANOVA and comparisons between bulk and sorted-infected cells for each virus are shown. **C.** Frequency of single nucleotide polymorphisms in HCoV-OC43-ORF4.5-mNG transcriptomic data, determined by lofreq, mapped to WT coordinates. The ORF4.5 insertion site is indicated by a dashed line.

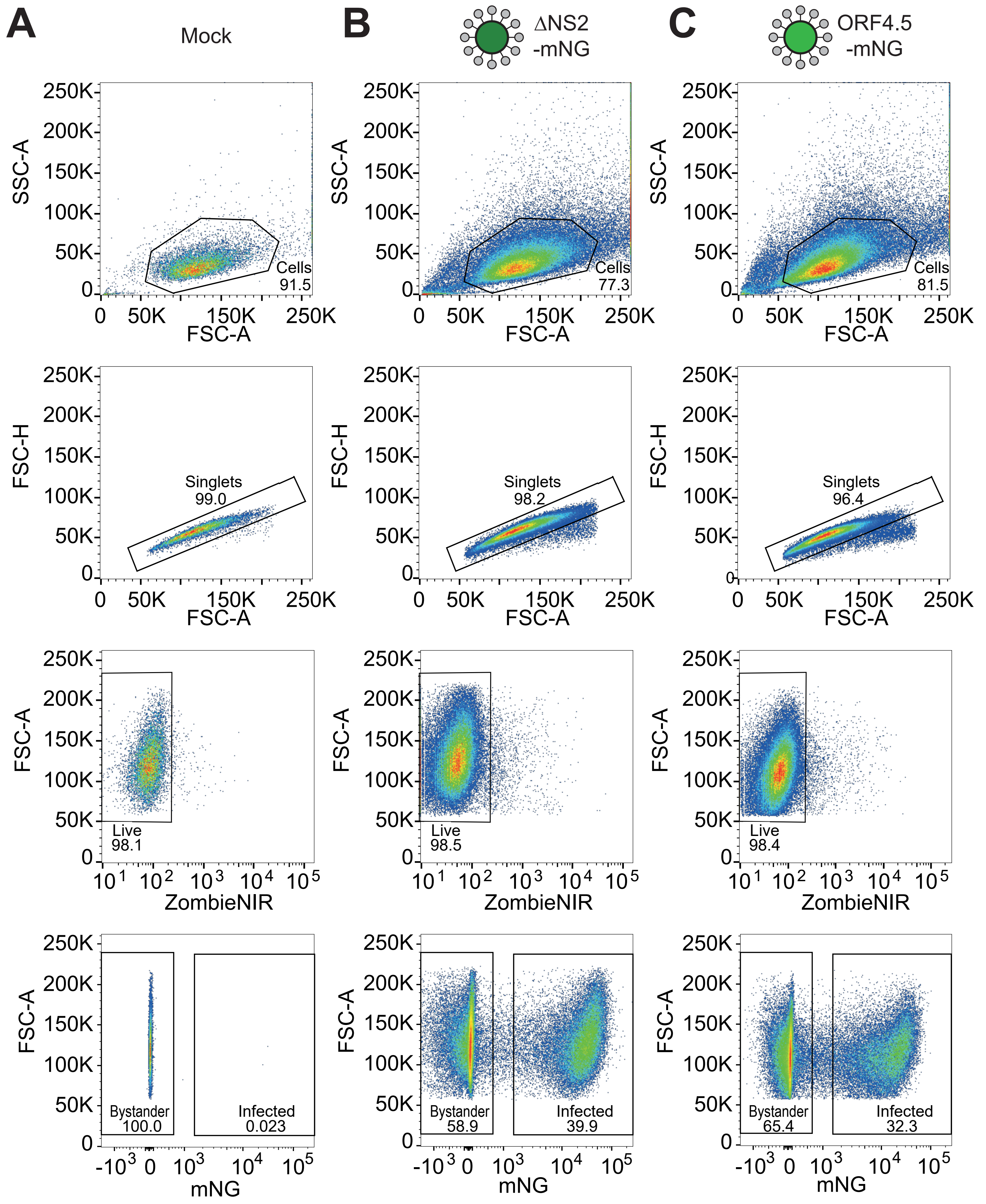


**Figure S6.** Sorting of cells infected with HCoV-OC43 reporter viruses. **A-C.** Gating strategy for isolation of live singlets and separation of infected (mNG-positive) and bystander (mNG-negative cells), for mock infected (A), HCoV-ΔNS2-mNG-infected (B) or HCoV-OC43-ORF4.5-mNG-infected (C) A549 cells, MOI 1, 24 hours post infection.

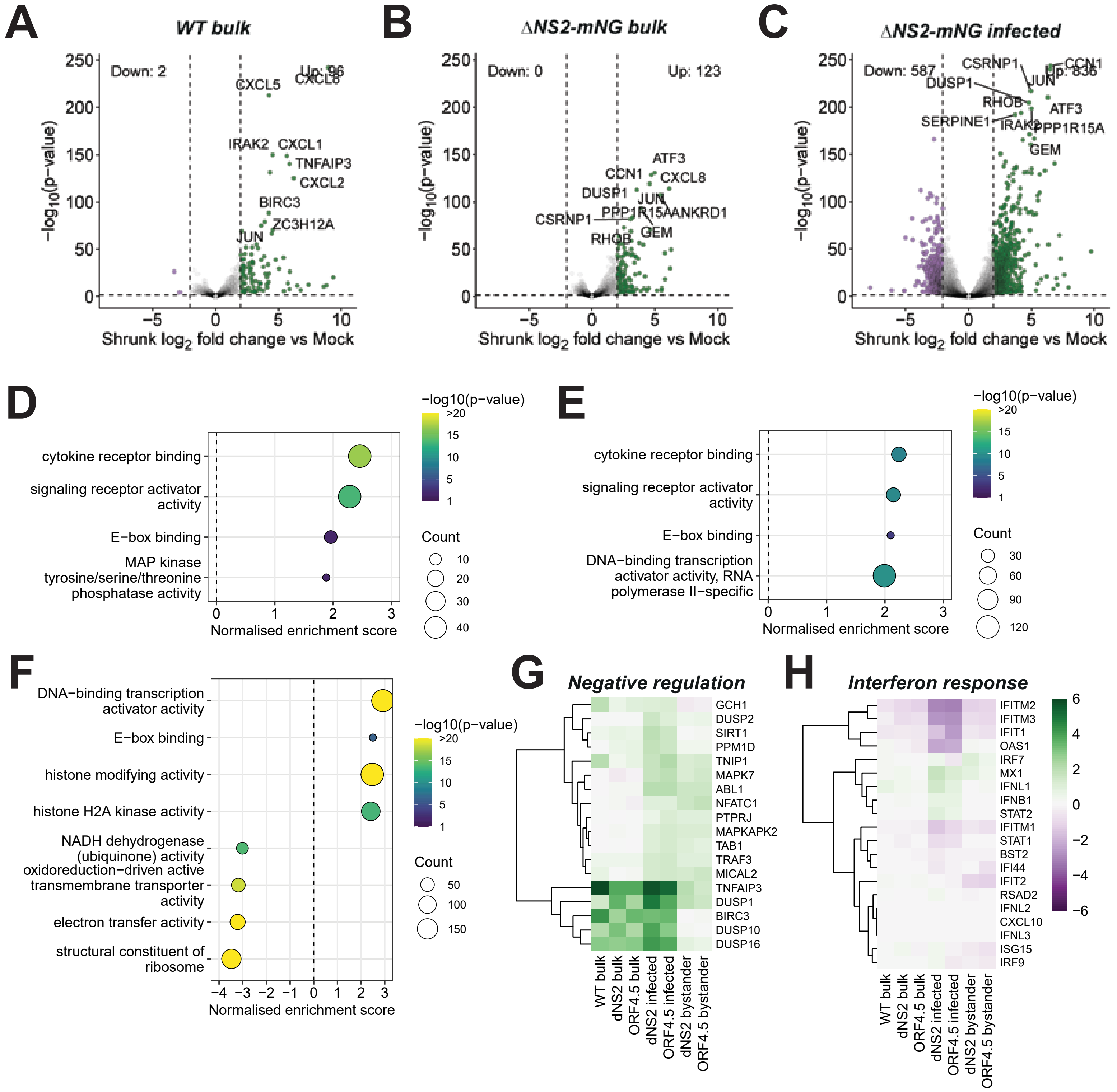


**Figure S7.** Host transcriptional responses in HCoV-OC43-WT and HCoV-OC43-ΔNS2-mNG infection. **A-C.** Volcano plots showing differentially expressed genes in HCoV-OC43-WT infection (A), or bulk (B) or sorted (C) HCoV-OC43-ΔNS2-infected cells, expressed as shrunk $\log_2$-fold change over mock infected cells. Significantly up- or downregulated genes (shrunk $\log_2$ fold change >±2, p value <0.05) are coloured and top 10 most significant genes are labelled. **D-F.** Corresponding gene set enrichment analysis of differentially expressed genes, based on molecular function. Functions are ranked by normalised enrichment score and coloured by significance. **G-H.** Heatmaps showing the expression of genes encoding negative regulators of NF-kappaB and MAPK signalling (G) and expression of classical type I IFN response genes (H), coloured by shrunk $\log_2$ fold change (shFC) in expression compared to mock cells.

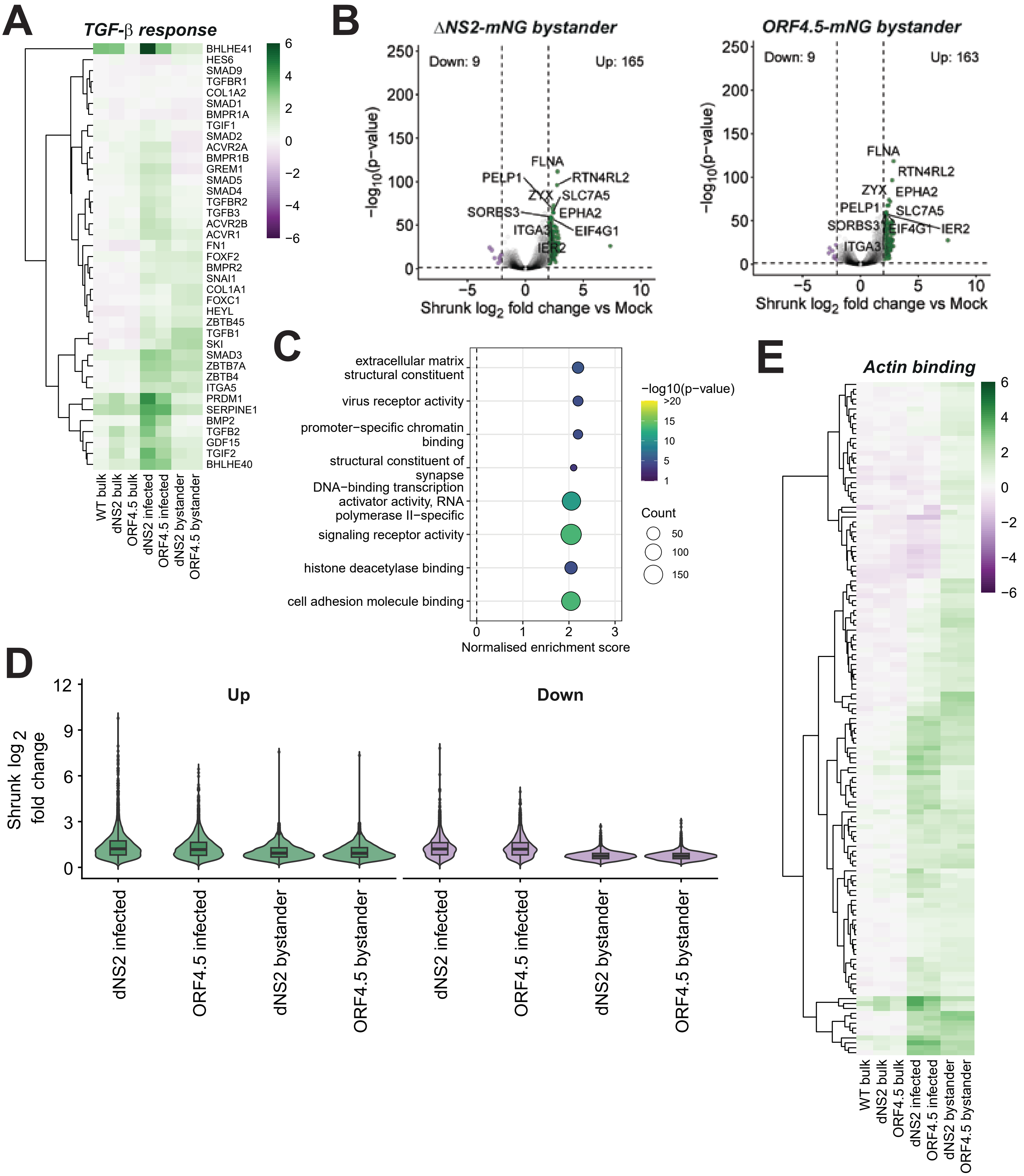


**Figure S8.** Actin filament induction in bystander cells. **A.** Heatmap showing the expression of genes involved in TGF-beta signalling, coloured by shrunk $\log_2$ fold change (shFC) in expression compared to mock cells. **B.** Volcano plots showing differentially expressed genes in bystander (mNG-negative) cells following HCoV-OC43-ΔNS2-mNG (left) or HCoV-OC43-ORF4.5-mNG (right) infection, expressed as shrunk $\log_2$-fold change over mock-infected cells. Significantly up- or downregulated genes (shrunk $\log_2$ fold change >±2, p value <0.05) are coloured and top 10 most significant genes are labelled. **C.** Gene set enrichment analysis of differentially expressed genes in ΔNS2-mNG bystander cells, based on molecular function. Functions are ranked by normalised enrichment score and coloured by significance. **D.** Summary of up- (green) and down- (purple) regulated gene expression in infected and bystander cells, shown as shrunk $\log_2$-fold change over mock-infected cells, for all genes where p<0.05. **E.** Heatmap showing the expression of genes related to actin binding, coloured by shrunk $\log_2$ fold change (shFC) over mock-infected cells.

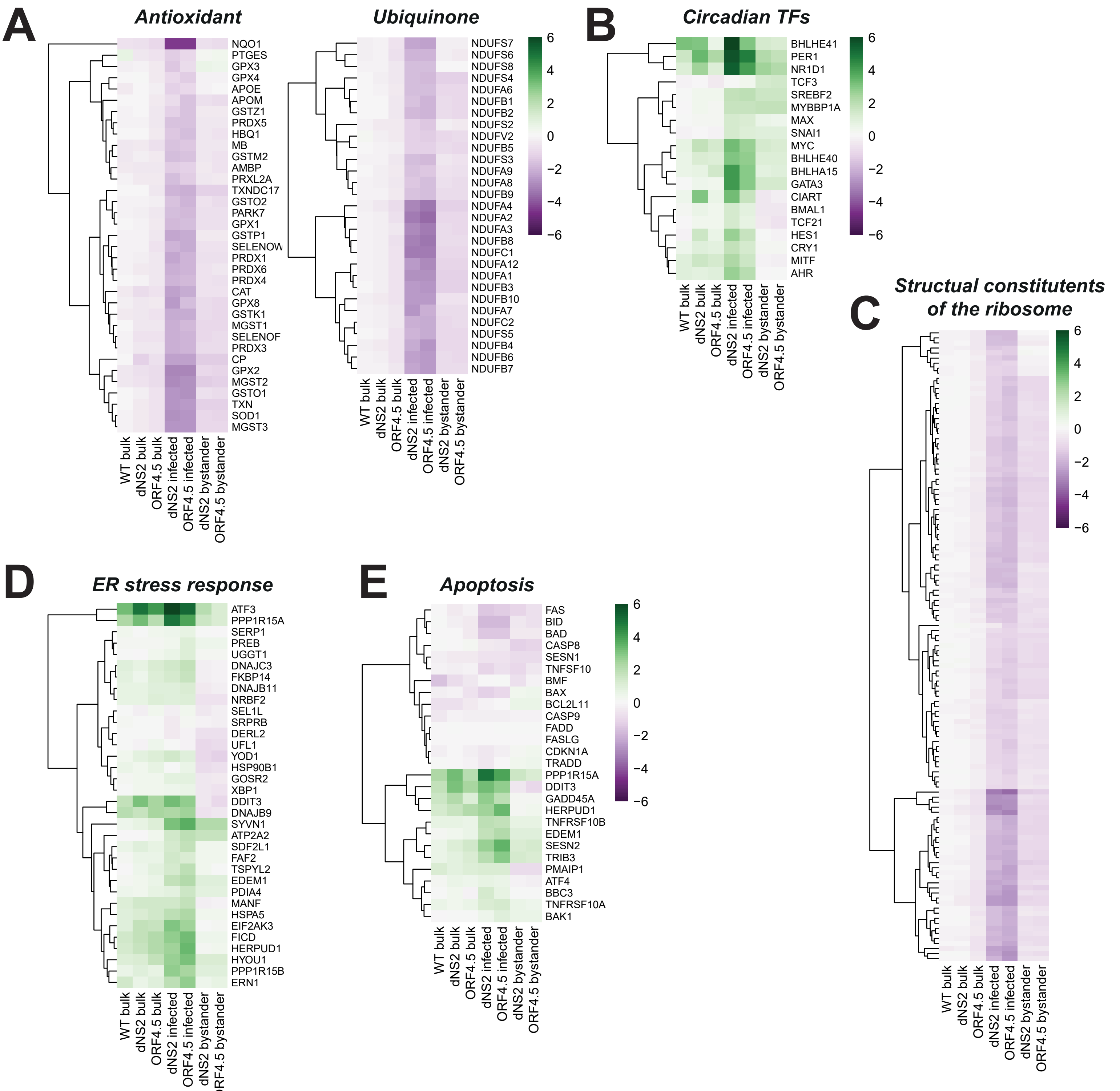


**Figure S9.** Downregulation of metabolism and induction of the ER stress response in infected cells. **A-E.** Heatmaps showing the expression of genes involved in oxidative phosphorylation (A), transcription factors associated with circadian rhythm regulation (B), structural constituents of the ribosome (C), the endoplasmic reticulum stress response (D) and pro-apoptotic markers (E), coloured by shrunk $\log_2$ fold change (shFC) over mock-infected cells. N.b. some ER stress response genes (D) are also represented in the apoptosis heatmap (E), since these pathways are interlinked.
